# Microsporidian *Nosema bombycis* secretes serine protease inhibitor to suppress host cell apoptosis *via* caspase BmICE

**DOI:** 10.1101/2024.06.27.600942

**Authors:** Maoshuang Ran, Jialing Bao, Boning li, Yulian Shi, Wenxin Yang, Xianzhi Meng, Jie Chen, Junhong Wei, Mengxian Long, Tian Li, Chunfeng Li, Guoqing Pan, Zeyang Zhou

**Author notes:** Corresponding author; (ZZ). These authors contributed equally to this work (Maoshuang Ran, Jialing Bao).

## Abstract

Microsporidia are a group of intracellular pathogens that actively manipulate host cell biological processes to facilitate their intracellular niche. Apoptosis is an important defense mechanism by which host cell control intracellular pathogens. Microsporidia modulating host cell apoptosis has been reported previously, however the molecular mechanism is not yet clear. In this report, we describe that the microsporidia *Nosema bombycis* inhibits apoptosis of *Bombyx mori* cells through a secreted protein NbSPN14, which is a serine protease inhibitor (Serpin). An immunofluorescent assay demonstrated that upon infection with *N. bombycis*, NbSPN14 was initially found in the *B. mori* cell cytoplasm and then became enriched in the host cell nucleus. Overexpression and RNA-interference (RNAi) of NbSPN14 in *B. mori’* embryo cells confirmed that NbSPN14 inhibited host cell apoptosis. Immunofluorescent and Co-IP assays verified the co-localization and interaction of NbSPN14 with the BmICE, the caspase 3 homolog in *B. mori*. Knocking out of BmICE or mutating the BmICE-interacting P1 site of NbSPN14, eliminated the localization of NbSPN14 into the host nucleus and prevented the apoptosis-inhibiting effect of NbSPN14, which also proved that the interaction between BmICE and NbSPN14 occurred in host cytoplasm and the NbSPN14 translocation into host cell nucleus is dependent on BmICE. These data elucidate that *N. bombycis* secretory protein NbSPN14 inhibits host cell apoptosis by directly inhibiting the caspase protease BmICE, which provides an important insight for understanding pathogen-host interactions and a potential therapeutic target for *N. bombycis* proliferation.

**Author Summary:** Microsporidia constitute a class of eukaryotic pathogens that exclusively reside within host cells. The species *Nosema bombycis* is the first microsporidian identified as the pathogen of silkworm Pébrine disease. In our research, we discovered how *N. bombycis* cleverly evades the host’s defenses. It has developed a strategy to survive inside host cells by manipulating host cell apoptosis, disarming the host cell’s self-destruct mechanism. In this study, we discovered that the *N. bombycis* secretes a serine protease inhibitor named NbSPN14, which infiltrates the cytoplasm of the host cell. The NbSPN14 interacts with the executioner Caspase protease BmICE within the silkworm’s apoptotic pathway, effectively neutralizing its apoptoic activity and thus curbing the apoptosis of the host cells.

## Introduction

Microsporidia are a large group of single-celled, eukaryotic, obligate intracellular pathogens which infect both vertebrates and invertebrate hosts [1, 2]. The first microsporidia species was identified in 1857, which described as the *Nosema bombycis* [3], as the causative pathogen of Pébrine disease in silkworms [4]. Since then, more than 1700 species in 220 genera have been identified [5]. The effects of microsporidia infections on their hosts cause significant economic loss to animal husbandry and are threats to public health [6, 7].

Apoptosis is a key feature of eukaryotic cells and is essential for the proper development of multicellular organisms [8]. In addition, as a defense mechanism, studies have demonstrated that host cells undergo apoptosis to purge invading pathogens,many pathogens, such as viruses and bacteria manipulate apoptosis to aid their intracellular survival [9–12]. The phenomenon of microsporidia modulating host cell apoptosis has been reported previously [13–18]. As early as 1999, Scalon et al. found that *Nosema algerae* (now called *Anncaliia algerae*) infected human lung fibroblasts cells (HLF) did not induce apoptosis, the survival time of infected cells *in vitro* increased several days compared to uninfected cells [19]. Aguila et al. have reported that *Encephalitozoon cuniculi* infection could suppress the apoptosis of host cells. Upon further analysis, it was discovered that the nuclear translocation of p53 and the activation of caspase 3 was markedly attenuated in Vero cells post-infection with *E. cuniculi* [18]. Similarly, both *Nosema apis* and *Nosema ceranae* reduce host cell apoptosis in bee epithelial cells [15, 16, 20, 21]. *N. bombycis* infecting the ovarian cells of Bombyx mori (BmN cells) has been shown to inhibit the host cell apoptosis by downregulating the expression of genes associated with the mitochondrial apoptosis pathway [17]. However, now which effector of microsporidia inhibit the host cell apoptosis is unknown.

Serine protease inhibitors (Serpins) are a group of protease inhibitors that are found in almost all organisms [26]. Serpins form complexes with target proteases, thereby, regulating various biological processes such as blood homeostasis [27], inflammatory responses [28], and cell apoptosis [29][30]. Serpins from pathogens have been reported to manipulate the host immune and apoptotic processes of host cells by regulating the activity of host proteases, which is beneficial for pathogen evasion against immune of host and pathogen self-proliferation [31]. SPI-2 and CrmA are the most extensively studied poxvirus serine protease inhibitors, they are non-essential for virus replication, but are involved in multiple immunomodulatory events [34]. It has been reported that SPI-2 and CrmA inhibit apoptosis and host inflammatory responses [35]. SPI-2 and CrmA target caspase 1 to protect virus-infected cells from TNF-mediated and Fas-mediated apoptosis as well as prevent the proteolytic activation of interleukin-1β [36, 37]. In the Microsporidia, a total of 53 serpins have been exclusively identified within the genus *Nosema*, and of these, 21 possess predicted signal peptides. Nineteen serpin members have been identified in *N. bombycis*, eight serpins with predicted signal peptides. The phylogenetic analysis demonstrated that the *N. bombycis* serpins clustered with the poxvirus serpins [32, 33].In general, above clues strongly suggest that the serpin protein encoded by *N. bombycis* may be involved in inhibiting host cell apoptosis.

Herein, we report that *N. bombycis* secretes a serpin protein, NbSPN14 into host cell cytoplasm, where it directly interacts with silkworm homolog Caspase 3, BmICE, the key executing effector in the silkworm apoptosis pathway. Subsequently, NbSPN14 inhibits the BmICE activity to suppress the host cell apoptosis, which ultimately facilitate the intracellular proliferation of *N. bombycis*.

## Results

### *N. bombycis* infection inhibits *B. mori* cell apoptosis

The presence of *N. bombycis* within infected host midgut tissues and culture cells was verified using an immunofluorescent assay by Hochest33342 nuclear staining (Fig. 1A). Terminal deoxynucleotidyl transferase dUTP nick end labeling (TUNEL) assay was then used to detect DNA fragmentation. Compared to un-infected and DNase I-treated positive controls, DNA fragmentations in the midgut tissues of infected silkworms were significantly reduced (Fig. 1B). The results from 1 to 7 days after infection were quantified by averaging the numbers of apoptotic cells from 5 to 10 observed fields under confocal microscopy. As shown in Fig. 1C, the percentage of midgut cell apoptosis was significantly decreased after *N. bombycis* infection. In addition, a silkworm embryonic cell line (BmE) was applied to confirm the above observations. The BmE cells were either un-infected, infected by *N. bombycis*, or further treated with the apoptosis inducer actinomycin D (Act D, 150 ng/mL). The results demonstrated that there was no significant difference in apoptosis between *N. bombycis*-infected and un-infected groups (Fig. 1D). However, when the two groups of cells were treated with actinomycin D, the percent of cell apoptosis of *N. bombycis*-infected BmE cells was significantly lower than that of the uninfected control (Fig.1E). Then, we determined the Caspase 3 activity in *N. bombycis*-infected cells, both before and after treatment with ActD. The findings revealed an elevation in Caspase 3 activity post-infection. However, upon ActD treatment, the Caspase 3 activity in the infected cells notably diminished compared to uninfected cells. (Fig.S1). The above results indicated that *N. bombycis* infection inhibits host cell apoptosis.

**Fig. 1.**
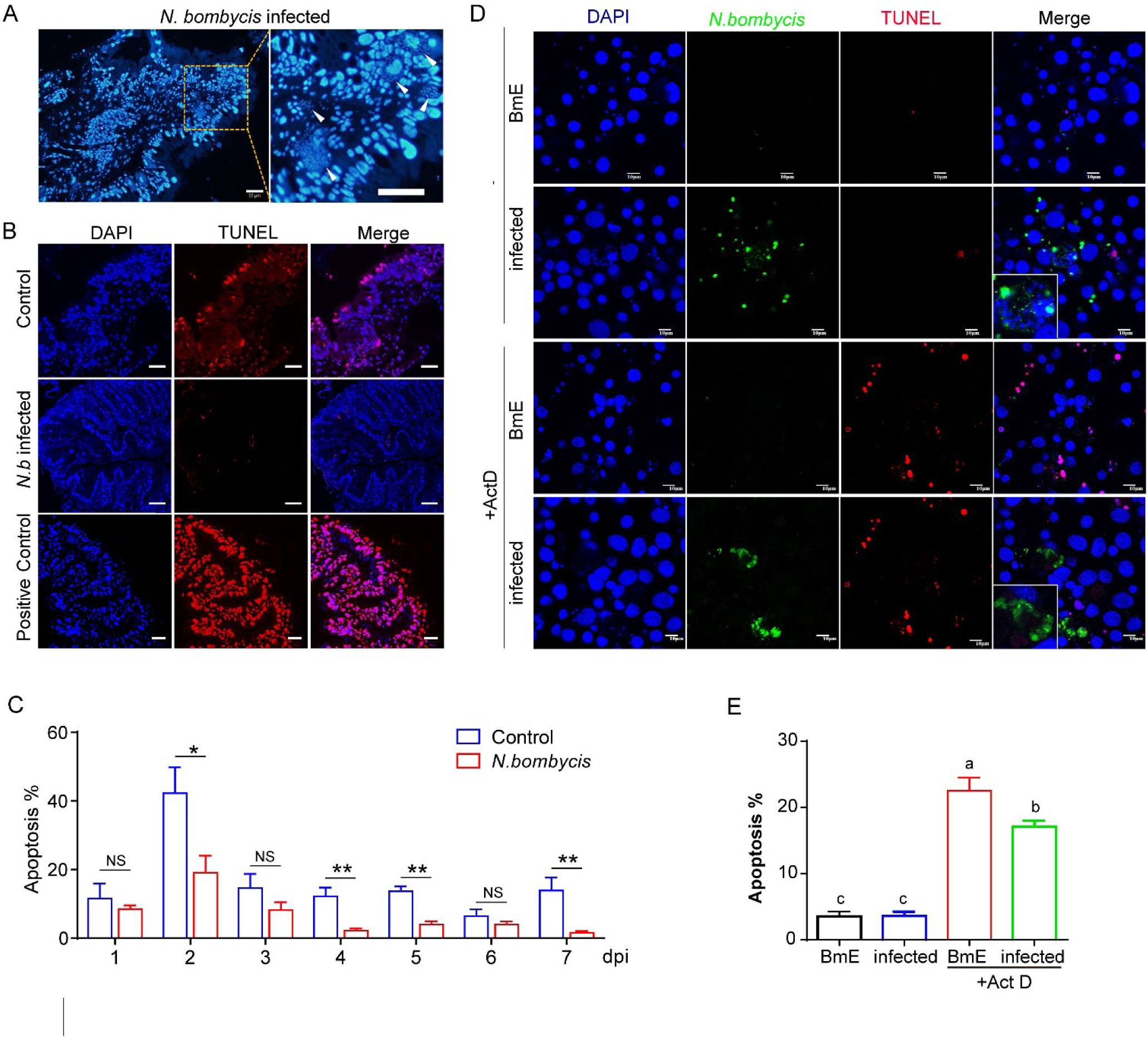
*N. bombycis* infection inhibits host cell apoptosis. A. Observation on the sections of silkworm midgut infected with *N. bombycis,* the white triangle is the *N. bombycis* nucleus marked by Hoechst33342, bar, 50 μm; B. TUNEL assay to detect apoptotic cells in transverse sections of infected and uninfected silkworm midgut, blue: nucleus, red: TUNEL positive signal; C. Quantitation of apoptosis in silkworm midgut cells 1 to 7 days after infection with *N. bombycis*. D. TUNEL analysis of the host cell apoptosis with *N. bombycis* infection 48h, green: *N. bombycis*; red: TUNEL positive signal; scale bar, 10 μm; E. Quantitation of apoptosis of BmE cells infected with *N. bombycis* for 48 hours, Different and same letters indicate values with statistically significant (p < 0.05) and non-significant (p > 0.05) differences, respectively.

### *N. bombycis* secretes NbSPN14 into its host cell

As a member of the *N. bombycis* serpin family, NbSPN14 was predicted with a secretory signal peptide (1-19Aa) and a typical serpin domain, the molecular weight is about 45kDa (Fig.2A). The polyclonal antibody of NbSPN14 reacts to the corresponding antigenic band in *N. bombycis-*infected cell lysates thus confirmed the presence of this protein (Fig.2B). Transcripts of NbSPN14 could be detected after infection, with the highest transcript level at 48 h post-infection (Fig.S2A). As shown in Fig.2C, *N. bombycis* was labeled by the specific antibody, while the NbSPN14 was found within host cells. Further analysis demonstrated that at 36 h after infection, NbSPN14 was mostly localized in the host cytoplasm, at 48-60 h post-infection, NbSPN14 had partially translocated into the host cell nucleus; and at 96 hours after infection (when the pathogenic load is very high) most of the NbSPN14 was translocated into the host cell nucleus. The transient expression of V5-tagged NbSPN14 in BmE cells also demonstrated that NbSPN14 could localize to the cytoplasm and nucleus of host cells (Fig.S2B). These findings confirm that NbSPN14 is a secreted protein, and the localization of NbSPN14 to the host cell nucleus suggests that NbSPN14 may interact with a host cell protein during proliferation.

**Fig. 2.**
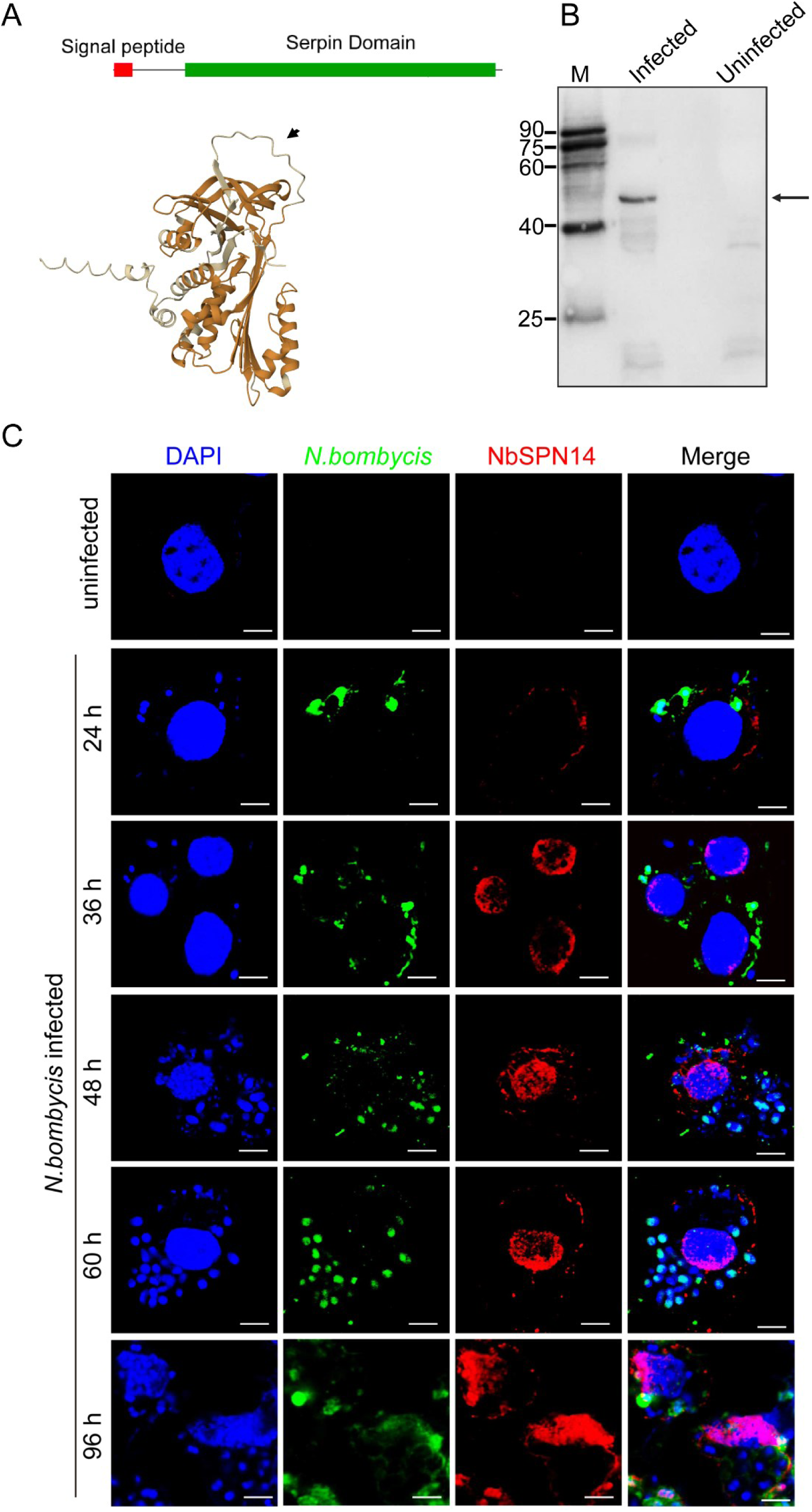
*N. bombycis* secretes NbSPN14 into host cell. A. NbSPN14 Signal Peptide, Serpin Domain and the predicted three-dimensional structure with typical structural characteristics of the serpin protein Reactive Center Loop exposure by AlphaFold Protein Structure Database. B. The protein expression level of NbSPN14 in infected BmE cell lines was analyzed using a Western blot, the arrow refers to the NbSPN14 band. C. Observations of the subcellular localization of NbSPN14 in infected BmE cells at 24, 36, 48, 60, 96 h post-infection by laser scanning confocal microscope. Nuclei are labeled with Hoechst 33342 (blue); NbSPN14 (red), and *N. bombycis* (green) labeled with relevant antibodies, respectively.

In order to study NbSPN14 function, NbSPN14 was expressed in cells to mimic the secretion of pathogens into host cells. The signal peptide sequence of NbSPN14 was removed from the coding sequence, and the remaining coding sequence was constructed into a pBac vector, then the recombinant vector was transfected into BmE cells (Fig.S3A). The transgenic cell line stably expressing NbSPN14 protein was obtained by Geneticin selection (Fig.S3B). PCR analysis shows that the NbSPN14 gene was integrated into the host cell genome and transcribed successfully. (Fig.S3C). Western blotting analysis showed that the cell line expressed the NbSPN14 protein successfully (Fig.S3D).

### NbSPN14 inhibits host cell apoptosis

CCK8 assays demonstrated that transgenic BmE cells expressing NbSPN14 possessed significantly higher cell viability compared to control cells (Fig.3A). Next, both groups of cells were treated with Act D, it was found that the proliferation activity of NbSPN14 transgenic cells were significantly higher than control ones (Fig.3B); TUNEL assay results showed that Act D could strongly induce apoptosis of BmE cells, and the cell number of apoptosis increased significantly after treatment with Act D; However, the apoptosis rate of the transgenic cell line expressed NbSPN14 was significantly lower than that of empty vector cell line (Fig.3C-D). Apoptosis can be visualized as a ladder pattern of 180-200 bp in standard agarose gel electrophoresis due to DNA cleavage by the activation of a nuclear endonuclease. The result showed the formation of the DNA ladder in gel electrophoresis by induction of apoptosis in NbSPN14 transgenic cells is much weaker than that of the control cell (Fig.3E). Similarly, the morphological changes of apoptosis can be clearly observed in phase with a larger number of apoptotic cells with the apoptotic body in the control group and slight apoptotic cells in the NbSPN14 transgenic cells (Fig.3F). What’s more, caspase 3 activity is an important indicator of apoptosis, as shown in Fig.3G, compared with control, the activity of Caspase 3 in the transgenic cell line expressed NbSPN14 was decreased significantly after Act D treatment, indicating that NbSPN14 play an important role in inhibiting host cell apoptosis.

**Fig. 3.**
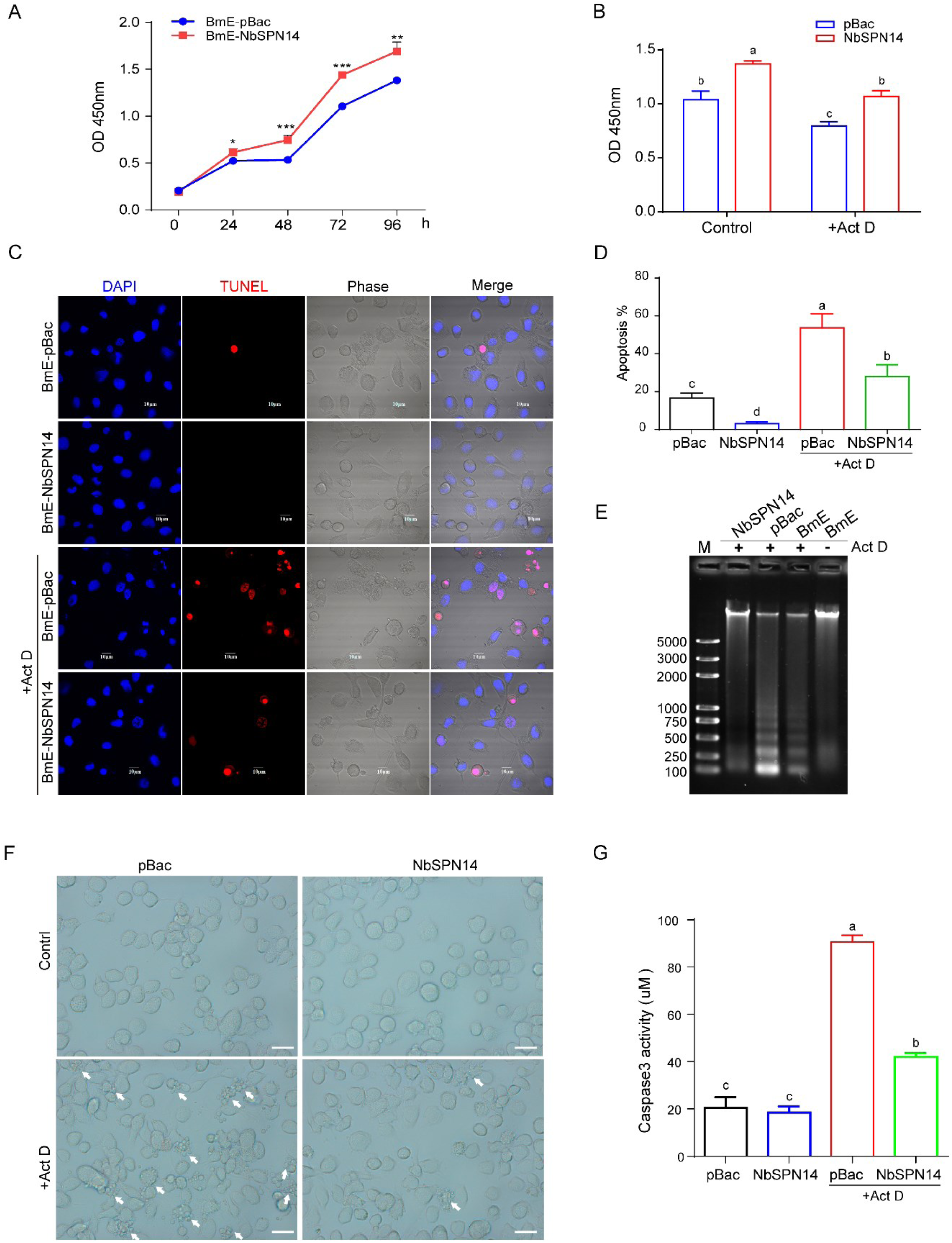
Inhibition of host cell apoptosis by overexpression of NbSPN14 in BmE cell lines. A. CCK-8 analysis of the cell proliferation after continuous culture of NbSPN14 transgenic cells at 0-96 h, and the x-coordinate was the continuous culture time; B. CCK-8 analysis of the NbSPN14 transgenic cells treated with Act D at 48h. C. TUNEL analysis the apoptosis of BmE-NbSPN14 transgenic cell line, red: TUNEL positive signal means the cell in a state of apoptosis; D. Quantification of the NbSPN14 transgenic cells apoptosis rate induced by Act D. E. DNA ladder analysis was used to detect the apoptosis of NbSPN14 transgenic cells after treatment with Act D. F. Morphological observation of NbSPN14 transgenic cells after induction of apoptosis by Act D treatment 12 h. Arrows indicate apoptotic cells, which are shrinking and bubbling to form many apoptotic vesicles. G. Caspase 3 activity analysis of NbSPN14 transgenic cells. Different and same letters indicate values with statistically significant (p < 0.05) and non-significant (p > 0.05) differences, respectively.

To further confirm the key function of NbSPN14 in inhibiting host cell apoptosis, we down-regulated the expression of NbSPN14 by transfecting the double-stranded dsRNA interference fragment targeting NbSPN14 gene into infected cells. The expression of NbSPN14 was detected by quantitative PCR and Western blotting, the results showed that the expression of NbSPN14 was down-regulated compared with the control group transfected with EGFP double-stranded dsRNA interference fragments (Fig.S4A-B). The results of caspase 3 activity assay showed that the infection of *N. bombycis* could inhibit the host cells’ Caspase 3 activity, down-regulating the expression of NbSPN14 led to the increase of caspase 3 activity of infected cells (Fig.4A). We utilized RNAi to block the expression of NbSPN14 in *N. bombycis* infected silkworm larvae. NbSPN14-dsRNA and the control EGFP-dsRNA were injected into the silkworms respectively after *N. bombycis* infection. The expression of NbSPN14 of the midgut determined by RT-PCR was drawn from 3 dpi to 4 dpi (Fig.S4C), and the results showed that NbSPN14 expression was successfully knocked down. TUNEL analysis result of midgut cell apoptosis after NbSPN14 interference was shown in Fig.4B. Compared with the uninfected group, the infected group midgut cells with the positive signals of TUNEL were significantly reduced. The apoptosis was significantly increased in response to the interference of NbSPN14 expression in the silkworm midgut compared with the infected group. These results confirmed that NbSPN14 is an important effector in *N. bombycis* inhibiting host cell apoptosis.

**Fig. 4.**
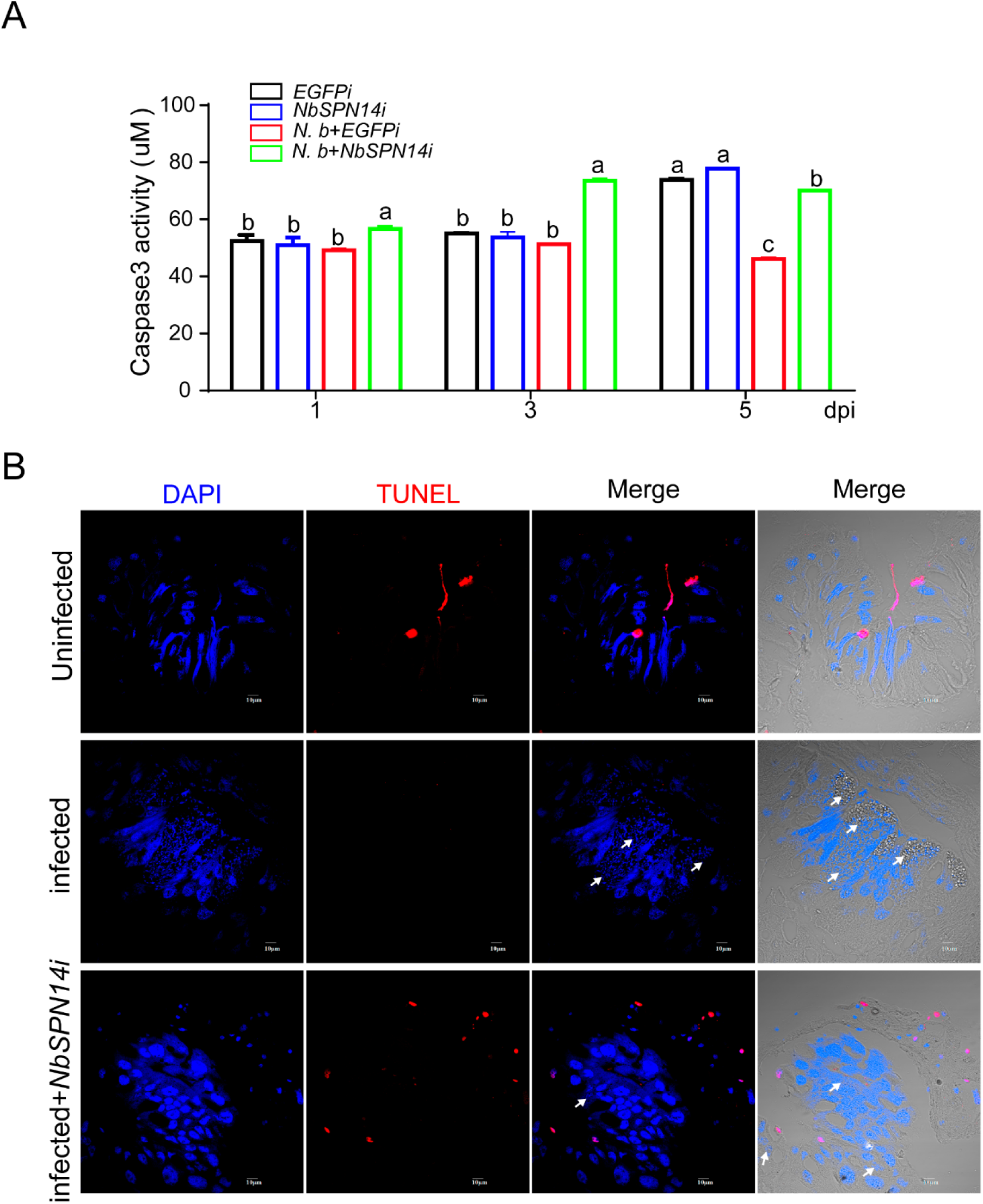
Host cell apoptosis increased after down-regulation of NbSPN14 expression by RNAi interference. A. Analysis of Caspase 3 activity in cells withNbSPN14-RNAi was significantly lower than the control group. Different and same letters indicate values with statistically significant (p < 0.05) and non-significant (p > 0.05) differences, respectively. B. The apoptosis of midgut cells of silkworm infected by *N. bombycis* were analyzed by TUNEL, the apoptotic signal of midgut cells increased significantly after NbSPN14 interference.

### NbSPN14 inhibiting apoptosis increases *N. bombycis* proliferation

To explore whether NbSPN14 inhibiting host cell apoptosis promotes the proliferation of pathogens in host cells, the proliferation of *N. bombycis* in host cells was analyzed following *N. bombycis* infection in NbSPN14 transgenic cell line and infected culture cells with NbSPN14 interference, respectively (Fig.5A). The copy number of *Nb β-tubulin* was used to the represent the pathogen load of the two groups. As shown in Fig.5B, the *Nb β-tubulin* copies in NbSPN14 transgenic cells were significantly higher than those in empty control cells at the middle and late stages of infection.However, the pathogen load in host cells was significantly reduced after NbSPN14 expression was down-regulated by RNA interference in infected cells at 5dpi (Fig.5C). The results showed that NbSPN14 inhibiting host cell apoptosis was important for *N. bombycis* intracellular proliferation.

**Fig. 5.**
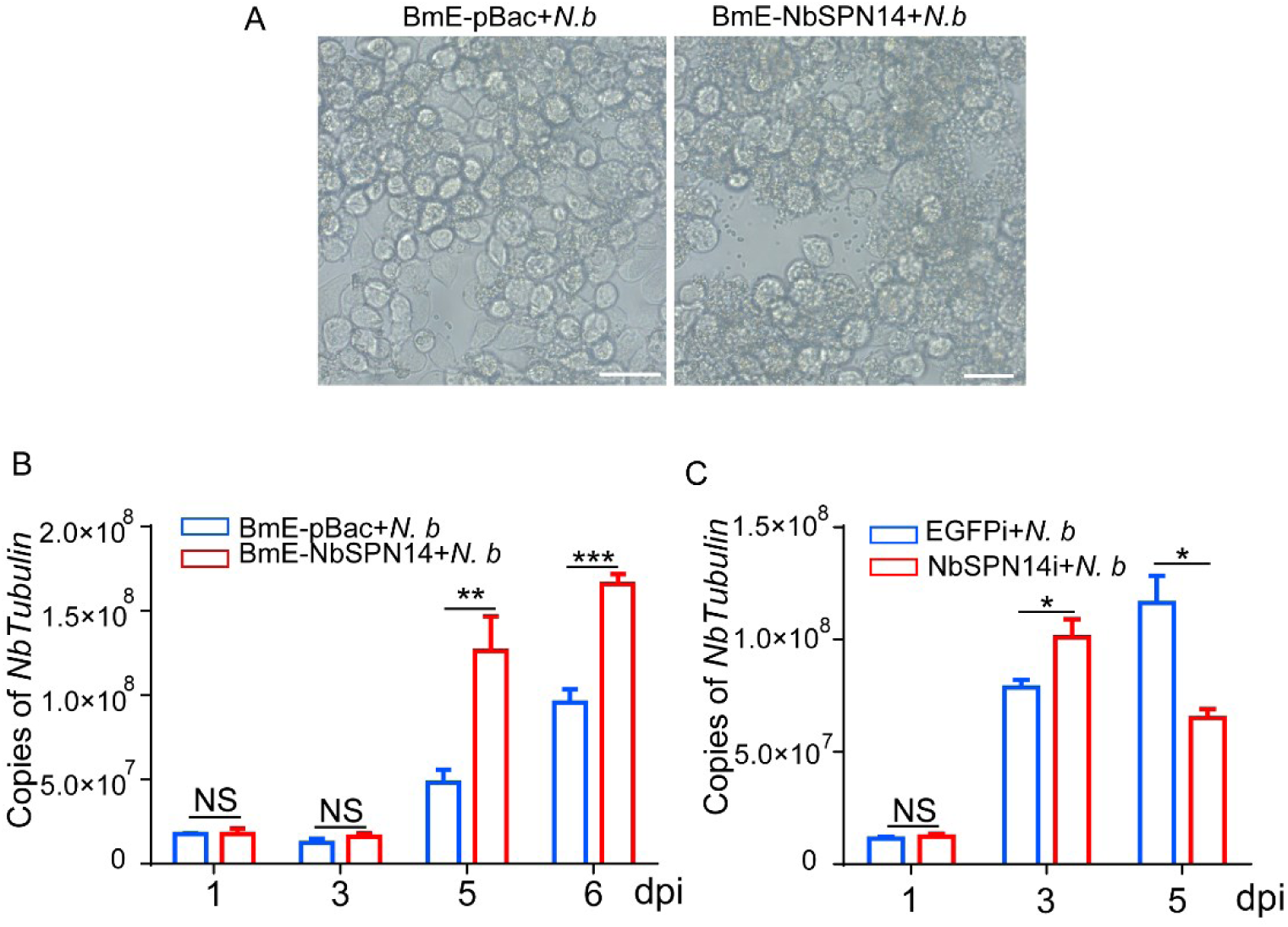
NbSPN14 can increase the proliferation of *N. bombycis* by inhibiting cell apoptosis. A. BmE-NbSPN14 transgenic cells morphology observation after 5 days of infection with *N. bombycis*, Bar scale 50 μm. B: The *β-tubulin* copies level in BmE-NbSPN14 transgenic cells was higher than that of empty Vector control cells in *N. bombycis* infection. C. The *β-tubulin* level in infected BmE cells knock-down NbSPN14. NS, no significant difference; *Indicates significant difference (P < 0.05), **indicates extremely significant difference (P < 0.01), ***indicates exorbitant difference (P < 0.001).

### BmICE is the target enzyme of NbSPN14

To investigate the target protein and the mechanism by which NbSPN14 inhibits the host cell apoptosis pathway, we analyzed Caspase 3 activity in cells following treatment with Ultraviolet light, Act D, and Staurosporine (STS). The results showed that the Caspase 3 activity of NbSPN14 transgenic cells after treatment with apoptosis inducers was significantly lower than that of the control group (Fig. S5A). NbSPN14 can inhibit apoptosis triggered by a variety of apoptotic agents, suggesting that the protein it targets plays a pivotal role in the apoptotic pathway. Then, we analyzed the transcription of genes related to the apoptosis pathway in NbSPN14 transgenic cells, and found that the pro-apoptotic gene *BmICE* was significantly up-regulated. In contrast, while the upstream apoptosis-related genes *BmDronc* and *BmDredd* showed no significant changes (Fig. S5B). Transcriptional analysis revealed a significant increase in BmICE levels; however, its activity was suppressed, which could be attributed to a rise in compensatory transcription. To ascertain whether NbSPN14 suppresses Caspase 3 activity by targeting upstream Caspase 9, we conducted enzyme activity assays. The results, depicted in Figure S5C-D, demonstrated that NbSPN14 is capable of inhibiting Caspase 3 activity instead of Caspase 9. Collectively, these findings imply that NbSPN14 may directly inhibit Caspase 3 activity.

The BmICE, the silkworm homolog of Caspase 3, behaving as the key executing effector in silkworm apoptosis pathway, is composed of the P10 and P20 subunits and includes the conserved pentapeptide motif QACRG. [22] (Fig.6A). It was predicted that the NbSPN14 P1 site at position 348th was aspartic acid D, which could be recognized by caspase. Furthermore, we discovered that the amino acid located at the P1 site of NbSPN14 is identical to the amino acid found at the P1 site of CrmA, a serpin protein encoded by the Cowpox virus. [37] Notably, CrmA is known to effectively hinders host cell apoptosis by suppressing caspase activity [43, 44]. This finding hints that NbSPN14 could possibly employ a similar mechanism to achieve inhibition of host cell apoptosis.

**Fig. 6.**
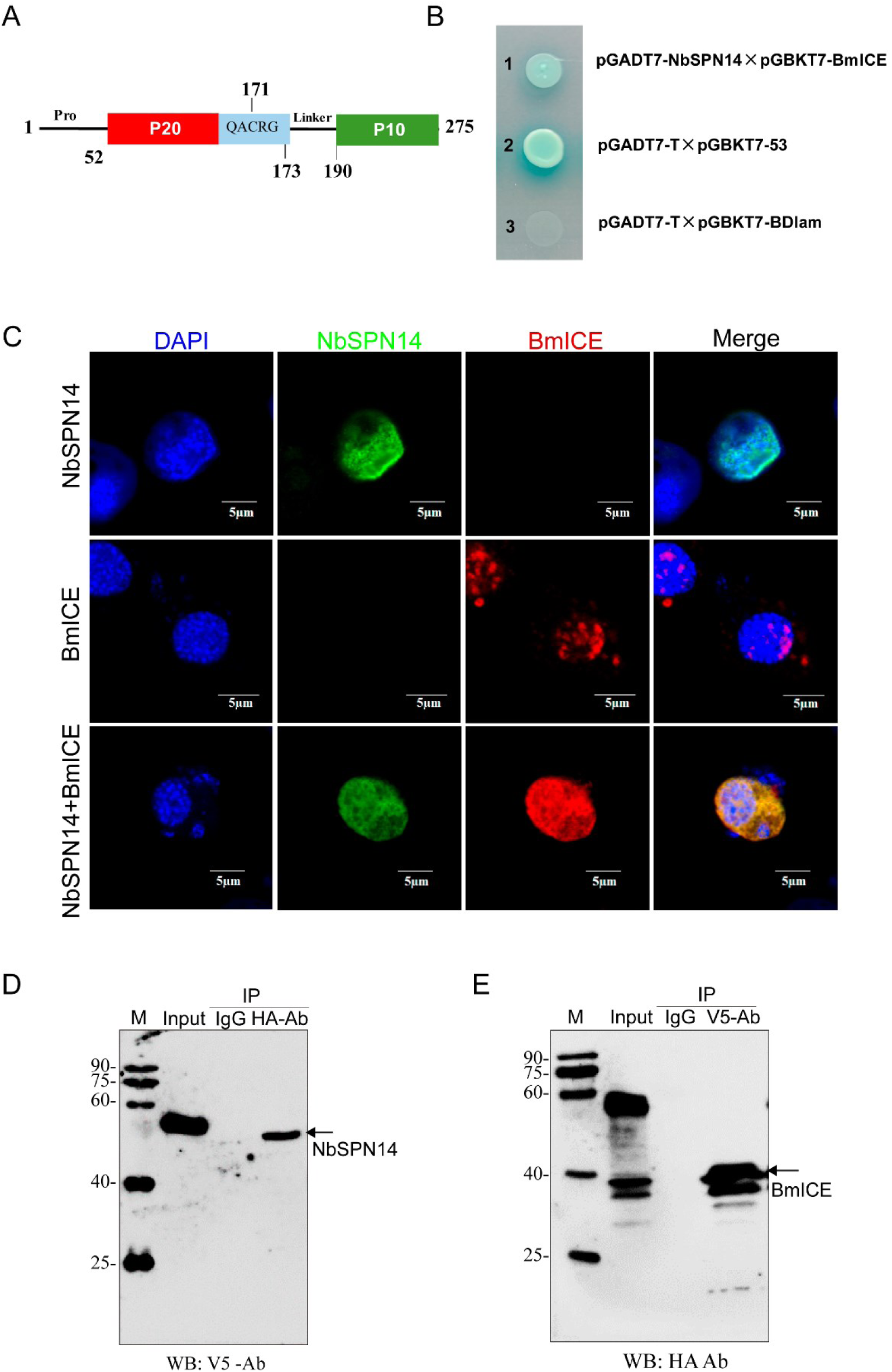
BmICE is the target protease inhibited by NbSPN14. A. The BmICE protein domain shows that BmICE has a pro domain and P10 and P20 subunits. B. Yeast two-hybrid assay to verify the interaction between NbSPN14 and BmICE. Interaction of pGADT7-NbSPN14 with pGBKT7-BmICE was screened by SD/-Ade/-His/-Leu/-Trp/X-α-gal/AbA medium. C. Co-localization assay of NbSPN14-V5 and BmICE-HA co-expression in BmE cells. Nuclei were stained by DAPI, NbSPN14 was detected using a V5 mouse monoclonal antibody and an Alexa-488 secondary antibody, BmICE was detected using an HA rabbit monoclonal antibody and an Alexa-594 secondary antibody. D. Lysates from cells co-expressing protein NbSPN14-V5 and protein BmICE-HA were subjected to Co-IP using an anti-HA antibody, followed by Western blot detection with an anti-V5 antibody. E. Lysates from cells co-expressing protein NbSPN14-V5 and protein BmICE-HA were subjected to Co-IP using an anti-V5 antibody, followed by Western blot detection with an anti-HA antibody. Input: samples of co-expressed cells, IgG: negative control.

Combined all the information above, we presumed that the candidate of the NbSPN14-inhibiting target protein was BmICE. To confirm the hypothesis, we utilized the yeast two-hybrid system, revealing a clear interaction between NbSPN14 and BmICE (Fig.6B). At the same time, we co-expressed NbSPN14 and BmICE in BmE cells, and found that they were co-localized in the host cytoplasm and nucleus, indicating that there may be an interaction between NbSPN14 and BmICE (Fig.6C). To further verify the interactions between NbSPN14 and BmICE, Co-IP was performed we simultaneously expressed fusion proteins: V5-tagged NbSPN14 and HA-tagged BmICE in BmE cells. The cell total protein samples were immunoprecipitated using a HA antibody, and the detection was performed using the V5 antibody. The V5 antibody confirmed the specificity of the NbSPN14 signal in immunoprecipitates, reverse verification can also be used to detect BmICE specific bands in immunoprecipitate as shown in Fig. 6D. The above results indicate that the NbSPN14 could interact with BmICE directly.

The amino acid residues of the P1 site in the RCL region of serpin protein play an important role in inhibiting target protease. Through site-directed mutagenesis, the 348th amino acid D of NbSPN14 was mutated to alanine A as NbSPN14^348A^, and the 347-349 amino acids were mutated into alanine A as NbSPN14^347-9AAA^(Fig.7A). Subcellular localization was analyzed after transient expression of NbSPN14 mutants in BmE cells. As shown in Fig.7B, NbSPN14 munants (including NbSPN14^348A^, NbSPN14^347-9AAA^) only localized in cytoplasm, could not enter the host cell nucleus. After constructing the cell line expressing NbSPN14 mutant protein, then we evaluated the function of the NbSPN14 mutant to inhibit host cell apoptosis (Fig.7C). The results showed that the Caspase 3 activity of mutated NbSPN14 transgenic cells was higher than that of wild-type NbSPN14 transgenic cells, which indicates that the P1 site amino acid residue D played a decisive role in NbSPN14 inhibiting the BmICE caspase activity.

**Fig. 7.**
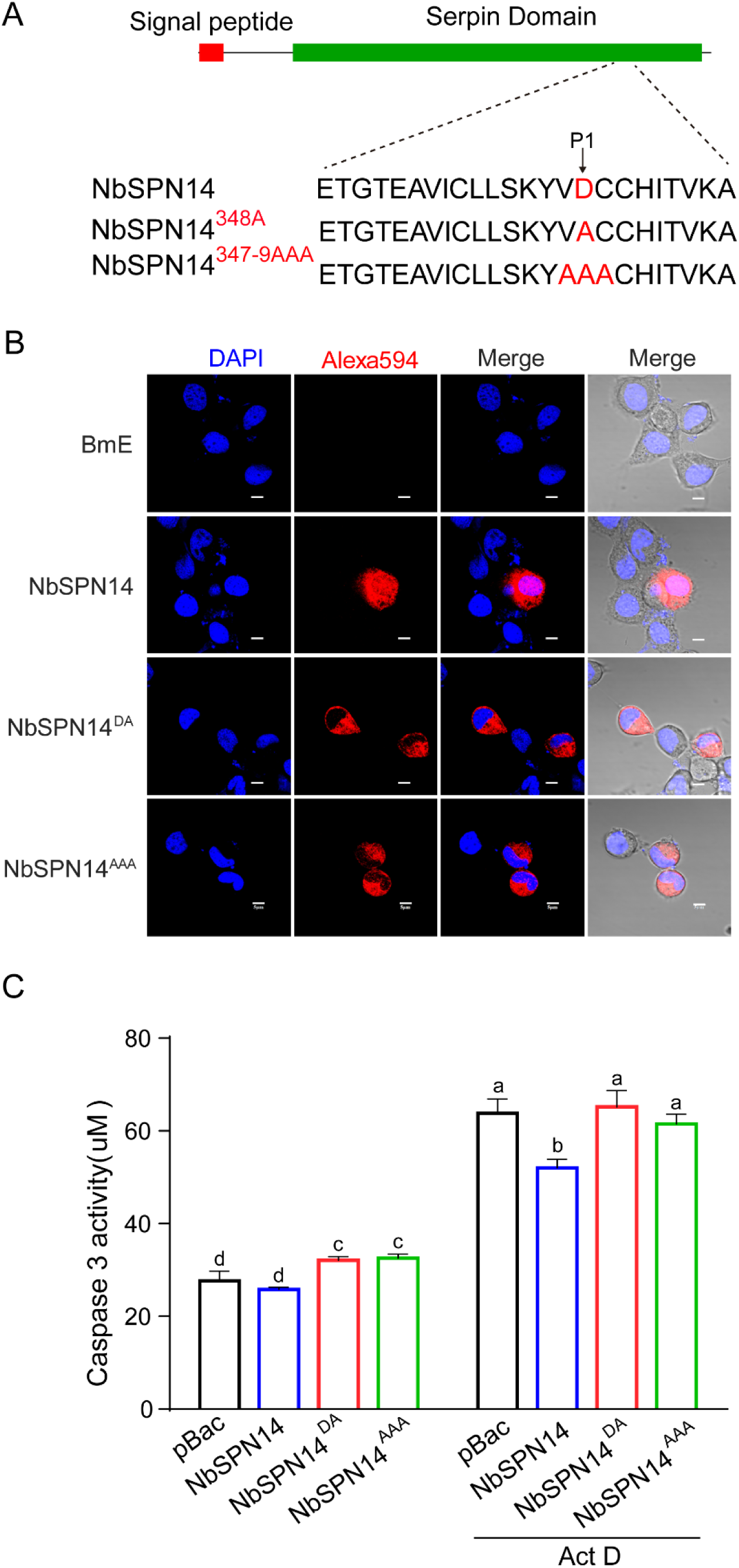
NbSPN14 P1 mutants are unable to enter the nucleus and inhibit apoptosis. A. NbSPN14 RCL sequence and P1 mutant schematic diagram. B. NbSPN14 and its mutant transient expression vector psL1180 [IE-NbSPN14mut-V5-SV40] were transfected into BmE cells for 48 hours to analyze the subcellular localization by immunofluorescence. The primary antibody uses the V5-tagged mouse monoclonal antibody, the secondary antibody was the Goat anti-Mouse IgG with Alexa Fluor® 594. C. Caspase3 activity analysis of NbSPN14 and NbSPN14 mutation transgenic cells with Act D treated 12h. Different and same letters indicate values with statistically significant (p < 0.05) and non-significant (p > 0.05) differences, respectively.

### BmICE plays an important role in silkworm apoptosis

To ascertain the role of BmICE in the apoptosis pathway of silkworm cells, we constructed a BmICE-overexpressing cell line and monitored the growth status of the cells. The results indicated that the growth index of BmICE-overexpressing cells was inferior to that of the control group, with an increased number of cells exhibiting apoptotic morphology and a significant rise in Caspase 3 activity. (Fig. 8A-B). Immunofluorescence assay (IFA) analysis revealed that BmICE, when overexpressed in cells, is localized to both the cytoplasm and the nucleus. Notably, during periods when cells display pronounced apoptotic features, including nuclear condensation, there is a marked accumulation of BmICE within the nucleus. (Fig. S6A).

**Fig.8.**
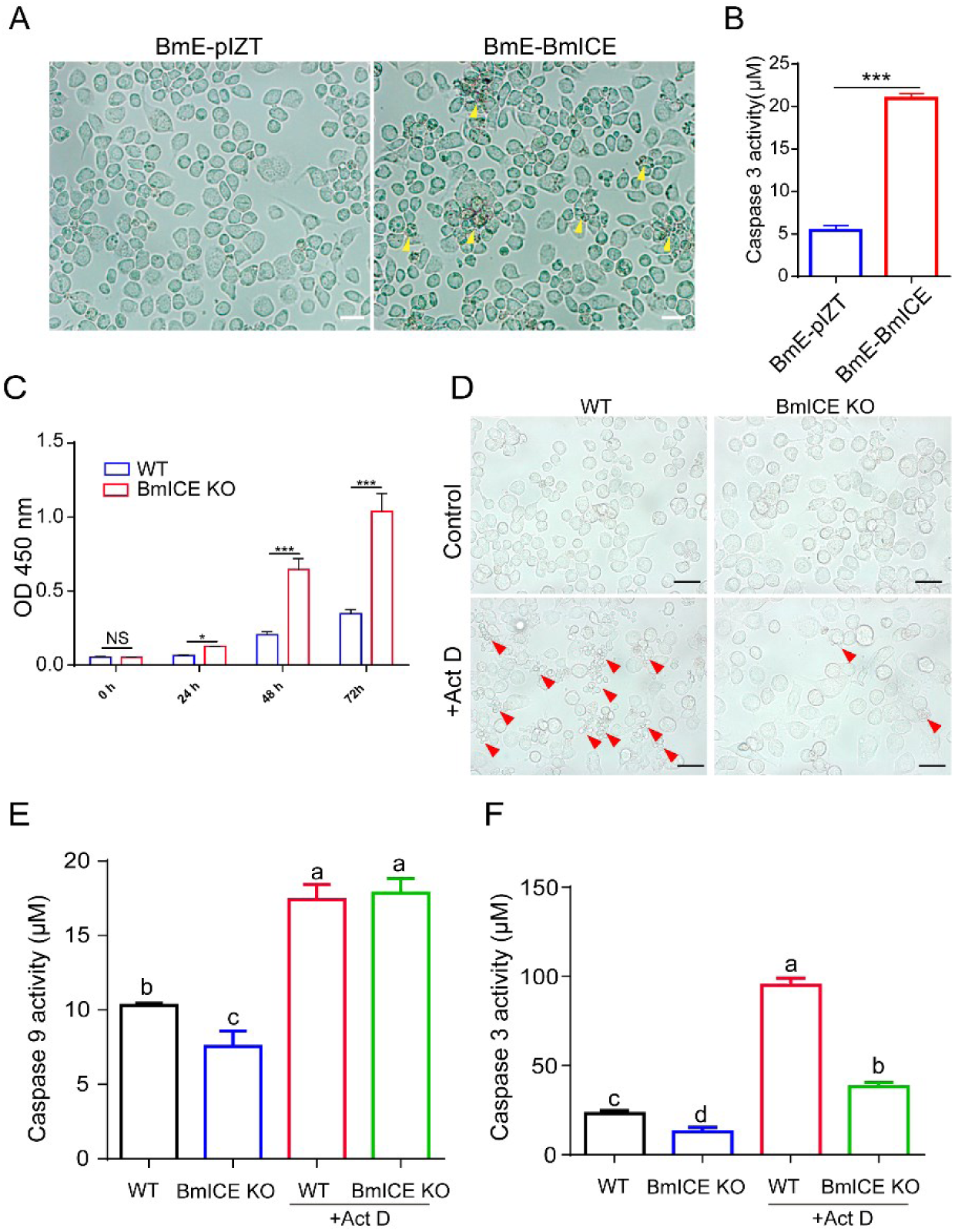
BmICE plays an important role in silkworm apoptosis. A. Observation of the morphology of BmICE overexpressed cells, white triangles indicate shrinking, foaming cells undergoing apoptosis; B: Caspase 3 activity of BmICE overexpressed cells were significantly higher than that of control empty vector cells. D. Observation of apoptotic morphology of BmICE knockout cells after Act D treatment 12h, the red triangle shows apoptotic cells. The apoptosis of BmICE knockout cells was significantly less than that of the control group. E. Analysis of Caspase 9 activity of BmICE knockout cells after Act D treatment showed no significant difference compared with the control group. F: Caspase3 activity of BmICE knockout cells after ActD treatment was significantly lower than that of control cells.

To further validate the function of BmICE in cell apoptosis. We used the CRISPR/Cas9 system to knock out the BmICE gene, and used double gRNA guidance to cleave the fourth and fifth exons of the BmICE gene and insert the GFP/Zeocin report/screened gene into its locus by homologous recombination (Fig. S6B). Primers were designed upstream and downstream of the recombination site for PCR validation, and the results showed that corresponding sized bands could be amplified, indicating successful knockout of the BmICE gene (Fig. S6C). BmICE knockout cells exhibited intact cell morphology and robust growth during the culture process. The proliferative activity of these cells was evaluated using the CCK-8 assay. Results showed that at 24h, 48h, and 72hpost-cell seeding, the cell viability of BmICE knockout cells was significantly higher than that of the empty vector control group (Figure 8C). Additionally, after a 12-hour treatment with Actinomycin D (Act D), the number of cells displaying apoptotic phenotypes in the BmICE knockout cell population was markedly lower than that in the control group (Figure 8D). Then, the Caspase 3 and Caspase 9 activities in BmICE knockout cells were detected and the results showed that the Caspase 3 activity of the knockout cells was significantly lower than that of the control group, there was no significant difference in Caspase 9 activity between the BmICE knockout cells and the control group (Fig. 8E-F). The results indicate that BmICE plays a crucial role in the apoptosis pathway of silkworm cells.

### NbSPN14 entering the host cell nucleus depends on BmICE

Both in *N. bombycis*-infected cells and overexpressed NbSPN14 in cells, it was found that NbSPN14 localized in the host cell cytoplasm and translocated into the host cell nucleus. However, subcellular localization prediction analysis showed that NbSPN14 has no nuclear localization motif, while BmICE can enter the nucleus[45]. We assume that NbSPN14 enters the nucleus after interacting with the target protein BmICE in the cytoplasm. To confirm our hypothesis, the transient expression vector of NbSPN14 was transfected into BmICE knockout cells, and the localization of NbSPN14 in BmICE knockout cells was analyzed by IFA (Fig.9). The results showed that the localization signal of NbSPN14 was distributed only in the cytoplasm, but not in nucleus in BmICE-knockout cells, while NbSPN14 could be localized throughout the entire cell including the nucleus in control cells. The results indicated that NbSPN14 relied on the interaction with BmICE to enter the nucleus.

**Fig. 9.**
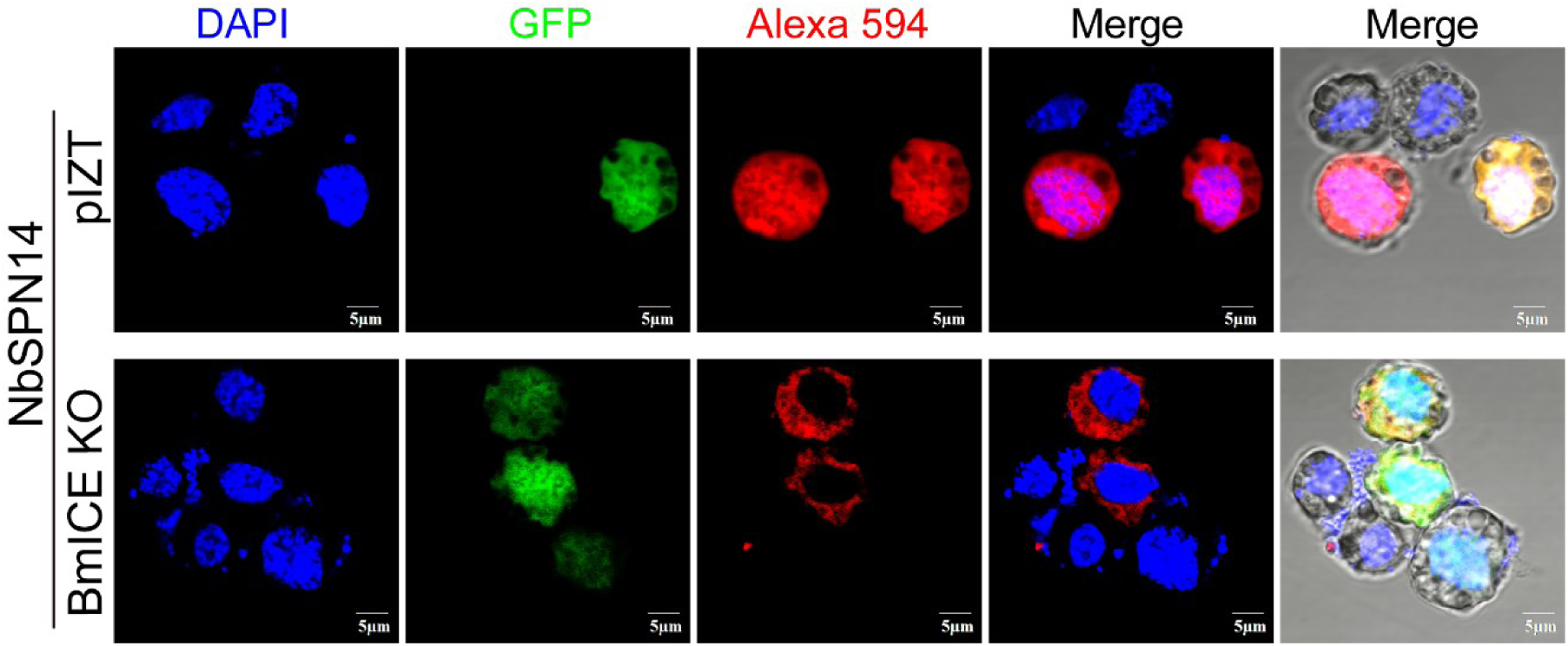
NbSPN14 entering the host cell nucleus depends on BmICE. A. Localization of transient expression of NbSPN14 in *BmICE* knockout cells. The recombinant NbSPN14 vector was transfected into *BmICE* knockout cells, and the location of recombinant protein NbSPN14-V5 in the cells was analyzed by V5 mouse monoclonal antibody with Alexa-594 conjugated secondary antibody. The results showed that NbSPN14 was not located in the host nucleus after BmICE knockout.

## Discussion

As a group of intracellular eukaryotic parasites, it was reported that microsporidian infection could inhibit host cell apoptosis [36–41]. However, it was unclear which effector was secreted into host cells to inhibit apoptosis. In this study, we demonstrated that *N. bombycis* secretes NbSPN14 into the host cell and inhibits the activity of the key apoptotic effector Caspase enzyme BmICE, thereby inhibiting host cell apoptosis. NbSPN14 binds to the activated BmICE in the cytoplasm and inhibits the caspase activity of BmICE; then the BmICE-Nbserpin14 complex enters the host cell nucleus. However, BmICE has been deactivated and lost its effector function--hydrolyzing cell components function in subsequent cell apoptosis, which thereby facilitates pathogen proliferation in the infected host cell (Fig.10).

**Fig. 10.**
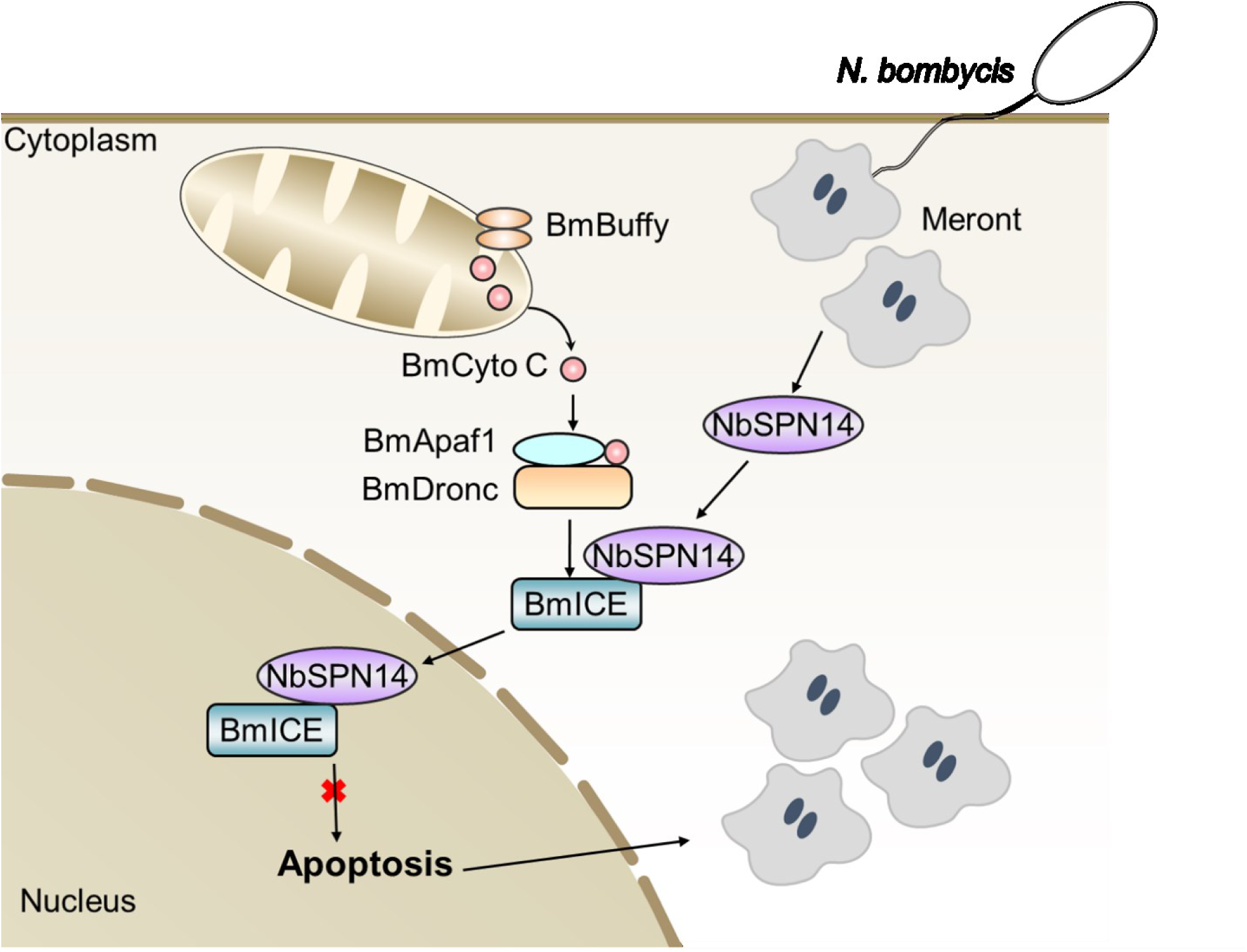
The schematic diagram of *N. bombycis* NbSPN14 inhibiting host cell apoptosis. *N. bombycis* infects the host cell through the polar tube injection and then enters become the meront that is the fission and proliferation stage. NbSPN14 is secreted into the cytoplasm of host cells by *N. bombycis* meront. Then, NbSPN14 binds to activated BmICE in the cytoplasm and subsequently translocated into the nucleus. NbSPN14 inhibits the BmICE activity, blocks apoptosis, and increases the *N. bombycis* proliferation.

Phylogeny analysis indicates that *N. bombycis* serpin genes are clustered with the poxvirus serpin genes [33]. The P1 site amino acid residue of the NbSPN14 reactive center loop was Aspartic acid, which is consistent with that of the Cowpox-Virus serpin, CrmA. It has been reported that CrmA inhibits apoptosis in a variety of cells [35]. Moreover, CrmA also inhibits the activity of Caspase protease, interleukin-1 beta-converting enzyme (ICE) in *Caenorhabditis elegans* [46]. Thus, we speculated that the target protein of NbSPN14 was the *B. mori* homolog BmICE. Our protein-protein interaction results confirmed that the target protein inhibited by NbSPN14 was BmICE. Although *N. bombycis* and poxvirus are far apart in the phylogenic tree of species, the molecular mechanism that *N. bombycis* secreting NbSPN14 to inhibit host cell apoptosis is similar to that of serpin in poxvirus [47].

Microsporidia, including *E. cuniculi, Vittaforma corneae, Nosema apis* and *Nosema ceranae* infection can inhibit host cell apoptosis [16, 18, 48]. However, the molecular mechanism(s) of inhibition of host cell apoptosis may not be identical in these microsporidia. The previous research results showed that microsporidia infection cloud inhibit host cell apoptosis through the typical Caspase pathway. The expression patterns of apoptosis-related genes were distinct after *E. cuniculi* and *V. corneae* infection of human macrophages THP-1, respectively; however, the Cysteine-aspartate protease (Caspase 3) activity was inhibited after THP-1 macrophages infected by both *E. cuniculi* and *V. corneae* [48]. Similarly, *E. cuniculi* infection prevented Caspase 3 cleavage in Vero cells [18]. Immunocytochemistry results demonstrated the depletion of Caspase 3 in the ventricular epithelial cells of honeybees infected with *N. ceranae* [20]. In this study, we confirmed that the secretion of NbSPN14 by *N. bombycis* inhibits caspase activity and inhibits host cell apoptosis. However, in the microsporidia, now only the *Nosema* genus has serpin family genes annotated, other microsporidia without serpin may inhibit host cell apoptosis through other effectors or different molecular mechanisms. Apoptosis Inhibitory Proteins (IAPs) are a highly conserved family of anti-apoptotic factors, which act directly on the caspase family and inhibit their activity [49]. IAP genes have been identified in a variety of microsporidia [50], so further study should focus on their functions involved in the regulation of host cell apoptosis. In addition, whether Serpins of other species of the *Nosema* genus are involved in inhibiting host cell apoptosis needs to be verified.

NbSPN14 is dependent on BmICE for entry into the host cell nucleus. NbSPN14 has a predicted secreted signal peptide but no nuclear localization signal (NLS). In the early stages of *N. bombycis* infection, NbSPN14 is secreted into the host cytoplasm, and then translocated to the host nucleus as the infection progresses (Fig 2). In the current study we have shown that NbSPN14 localizes to the host cell cytoplasm and interacts with BmICE. The interaction leads to the loss of caspase activity and the inhibition of host cell apoptosis. A previous study has shown that the Caspase 3 homolog BmICE acts as an effector factor in regulating apoptosis [22, 51]. Caspase 3 is predominantly localized in the cytoplasm in the form of pro-enzyme [45]. During the process of apoptosis, activated Caspase 3 is translocated into the nucleus to cleave its nuclear substrates, resulting in characteristic apoptotic nuclear changes such as DNA fragmentation, chromatin agglutination, and nuclear disruption [52, 53]. We demonstrated that NbSPN14 binds to BmICE in the host cytoplasm and then was translocated to the host cell nucleus. We expressed NbSPN14 in BmICE knockout cells and found that NbSPN14 could not be translocated into the host cell nucleus, confirming that NbSPN14 inhibits BmICE activity in the cytoplasm and then be translocated into the host cell nucleus along with BmICE.

NbSPN14 inhibits host cell apoptosis, which increases pathogen proliferation. Intracellular infection depends on the survival of host cells for pathogen proliferation [54]. Inhibition of host cell apoptosis by pathogenic secreted effectors is a common strategy for the survival of intracellular pathogens [55]. Herein, we demonstrated that NbSPN14 is a key effector molecule in host cell apoptosis inhibition, which is essential for robust *N. bombycis* proliferation. It raises the possibility that therapeutic compounds targeting this pathway or genetic manipulation of the host could protect silkworms from infection with *N. bombycis*. Previous studies have shown that the transgenic expression of monoclonal antibody single-chain variable fragments (scFvs) targeting *N. bombycis* protein in silkworm or host cell lines reduced *N. bombycis* infection and proliferation [56–58]. As host cell apoptosis promotes the clearing of infected cells and reduces the load of *N. bombycis*, we can express monoclonal antibody scFv against NbSPN14 in silkworm to block the *N. bombycis* serpin inhibiting host cell apoptosis; as a result, the resistance of the host silkworm to *N. bombycis* will enhance. In addition, serpin usually contains a conformation relaxed reactive center loop (RCL) that is cut between P1 and P1’ residues by a target protease [59, 60]. The RCL P1 site amino acid residues are critical for serpin function and specificity [61]. After the reaction between the protease and serpin, the protease is trapped in a covalent complex with serpin[62]. Consequently the RCL is rapidly incorporated as a new central β-strand into the serpin A β-sheet[63]. It has been reported that specific RCL-derived peptides could mimic the RCL insertion strand within serpin domain 4A [64–67], and cause the serpin function loss [68]. Thus, it is feasible to devise peptides derived from NbSPN14 RCL that can hinder the function of NbSPN14, thereby preventing the inhibition of host cell apoptosis, ultimately aiming to bolster the silkworm’s resistance against Pébrine disease.

## Materials and methods

### Silkworm rearing and cell culture

The silkworm strain Dazao was obtained from the Gene Resource Library of Domesticated Silkworm (Southwest University, Chongqing, China). Silkworms were reared with fresh mulberry leaves at a temperature of 25°C and relative humidity of 70%. Silkworm cell lines were maintained at 28 °C in Grace’s medium (Thermo Fisher Scientific, United States) supplemented with 10% (V/V) fetal bovine serum (FBS) (Thermo Fisher Scientific, United States) and 1% (V/V) penicillin-streptomycin[38].

### *N. bombycis* infections *in vitro* and *in vivo*

Mature *N. bombycis* spores were isolated from infected silkworm pupae and purified by Percoll density gradient centrifugation (21,000g, 40 min) [39]. Purified *N. bombycis* mature spores were treated with 0.1 M KOH 3 minutes and then the spores were incubated with BmE cells at 10:1 ratio in Grace’s medium at 28°C. The medium was replaced after 2 hours, and the cells were cultured at 28 °C in Grace’s medium supplemented with 10% (V/V) fetal bovine serum (FBS) and 1% (V/V) penicillin-streptomycin. Silkworms were raised to the third day of the fourth instar, and then were fed on mulberry leaves (1 cm^2^) spread with 105 mature spores.

### Tissue paraffin section

For histological analysis of midgut cell apoptosis (infected and uninfected), the midgut of the fourth abdomen was isolated and fixed with 4% paraformaldehyde for 24h at room temperature, then dehydrated through the series of ethanol, and embedded in paraffin. Then this material was sectioned into 5-μm slices and placed on slides. After deparaffinisation and hydration, sections of the slides were stained with hoechst33342 and TUNEL to analyze the apoptosis of midgut cells.

### Terminal uridine nick-end labeling (TUNEL) assay

Apoptosis in Silkworm midgut and cultured BmE cells was detected using a terminal deoxynucleotidyl transferase-mediated dUTP-biotin nick end labeling (TUNEL) assay (Beyotime, China).

For this assay, the midguts of infected silkworms were fixed in 4% paraformaldehyde, embedded in paraffin for 24h, sectioned into 5-μm slices, and placed on slides. After deparaffinisation and hydration, the slides were stained with TUNEL by incubation with TdT Reaction Buffer at 37 °C for 10 min, followed by the incubation with a TdT reaction mixture at 37 °C for 1 h. The slides were then incubated with a TUNEL reaction cocktail at 37 °C for 30 min, counterstained with Hoechst 33342. Images were acquired using a confocal microscope (OLYMPUS).

For this assay, BmE cells were cultured in twelve well plates and infected with 1×10^6^ spores for 48h. Cells were fixed with 4% paraformaldehyde for 15 min then incubated with PBS containing 0.3% Triton X-100 incubated for 5 minutes at room temperature. The cells were then stained with TUNEL and IFA. Images were acquired using a confocal microscope (OLYMPUS).

### Indirect Immunofluorescence Assay (IFA)

In order to characterize the location of NbSPN14 in the process of infection. BmE cells infected for varying lengths of time with *N. bombycis* were fixed with 4% paraformaldehyde for 10 min at room temperature, washed three times with 1xPBS, and then permeabilized using 0.1% Triton X-100 for 15 min. The cells were then blocked in 1xPBST containing 5% BSA and 10% goat serum for 1 h at room temperature. Next, the cells were incubated with mouse and rabbit polyclonal antibodies against NbSPN14 (anti-NbSPN14) and *N. bombycis* (anti-*N. b* total protein) diluted 1:100 in a blocking solution for 2 h at room temperature. The cells were then washed three times with 1xPBST, and incubated for 1 h with a 1:1000 dilution of Alexa Fluor 488 conjugate Goat anti-Mouse IgG (Invitrogen A32723, Rockford, Illinois, USA) and Alexa Fluor 594 conjugate Goat anti-Mouse IgG (Invitrogen A32742, Rockford, Illinois, USA) in a dark moist chamber at room temperature. The cell nucleus was stained with DAPI (1:1000 dilution, Sigma-Aldrich 28718-90-3, St. Louis, MO, USA) at room temperature for 15 min. The samples were finally observed and photographed using an Olympus FV1200 laser scanning confocal microscope.

### Protein structure prediction

The protein structure of NbSPN14 was modeled using AlphaFold2 online (https://alphafold.com/) employing the default settings. [40] (no structural template was used). All five AlphaFold2 models were then tested. The structures of the five AlphaFold2 models were consistent and the first model was selected for the predicted structure of NbSPN14.

### Western blot

Western blotting was used to detect the expression of NbSPN14 in infected cells, the total protein of BmE cells infected with *N. bombycis* was extracted using 300 μl RIAP strong lysis buffer containing PMSF ice for 15 min, and then centrifuged at 12,000 g at 4°C for 15 min. The supernatant was then used for 10% SDS-PAGE and then transferred to a PVDF membrane (Roche). The PVDF membrane was then blocked with 5% (w/v) skim milk (37 ℃, 1h), followed by incubation with the antibody for 1.5 h at room temperature. Membranes were then washed 3 times (5 min each) with TBST buffer (10 mM Tris, 150 mM NaCl, 0.1% Tween-20), incubated with goat anti-mouse/rabbit IgG peroxidase antibody for 50 min at room temperature, and washed 3 times with TBST buffer. The antibody binding was detected using Clarity™ Western ECL substrate (Bio-Rad). Western blot experiments, such as detecting protein expression in transgenic cells and verifying interactions used the same methods as described above.

### Construction of NbSPN14 transgenic cell lines

The amino acid sequence of NbSPN14 was submitted to SignalP 6.0 Server (http://www.cbs.dtu.dk/services/SignalP/) and NCBI (https://www.ncbi.nlm.nih.gov/) for the signal peptide and domain predictions. The *NbSPN14* of *N. bombycis* (GenBank Accession No. FJ705061.1) was amplified from genomic DNA (gDNA) by PCR. The NbSPN14 signal peptide sequence was removed, and the forward primer containing a BamH I restriction site and the reverse primer 5 containing a Not I restriction site were used. The amplification reaction consisted of 30 cycles of 95 °C for 15 s, 55 °C for 30 s, and 72 °C for 1 min. The PCR products were recovered, integrated into the pSL1180[IE2--SV40] vector, then the [IE2-NbSPN14-SV40] sequence fragment was amplified by primer pairs ‘pBac-F - pBac-R’ and ligated into the PiggyBac vector [A3-EGFP-SV40+A3-Neo-SV40-]. The recombinant vector PiggyBac [A3-EGFP-SV40+A3-Neo-SV40+IE2-NbSPN14-SV40] was extracted from the DH5α cells with the TIANpure Mini Plasmid Kit (TIANGEN, China). BmE cells were transfected with 2 μg of this recombinant plasmid and A3 helper plasmid using X-tremeGENE HP DNA Transfection Reagent (Roche, Switzerland), and the culture medium was changed after 6 h. Three days later, the cells were cultured in Grace insect complete medium containing G418 (200 μg/mL) (Merck, Germany), and the culture medium was changed once every 3 d. Screening continued for two months until the proportion of cells with fluorescent green exceeded 98% [41].

### CCK8 assay

The cell proliferation of BmE cells was tested using cell counting kit-8 (CCK8) (MCE, HY-K0301). According to the instructions of manufacturer, BmE cells in the logarithmic growth phase were seeded into the 96-well plates (5000 cells/well). Cells were incubated at 280c for 0, 24, 48, 72, or 96 h and 10 μL of CCK8 reagent was added to each well After 3 h of incubation, the absorbance at 450 nm was measured to determine the cell viability using a multifunctional enzyme-labeling instrument. All experiments were performed in triplicate.

### Caspase 3/9 activity assay

NbSPN14 Transgenic cells were seeded into a 6-well culture plate and incubated in Grace supplemented with 10% FBS. After being treated as described above, protein extracts were prepared following the manufacturer’s instructions using a Bradford Protein Assay kit (Beyotime Institute of Biotechnology, Nantong, Jiangsu, China). Caspase 3 activity was measured using a Caspase 3 Activity Assay kit (Beyotime Institute of Biotechnology, Jiangsu, China) in which cell extracts were mixed with Ac-DEVD-*p*NA substrate for 2 h at 37 °C in 96-well plates prior to colorimetric measurement of *p*-nitroanilide product at 405 nm. The Caspase 9 activity assay was the same as the Caspase 3 activity assay, except that Ac-LEHD-*p*NA is used as the substrate.

### RNA interference

The sequence of *NbSPN14* was submitted to BLOCK-iT™ RNAi Designer (http://rnaidesigner.thermofisher.com/rnaiexpress/design.do). Two fragments that contain five potential interferential dsRNA fragments were amplified by the primers NbSPN14-1T7 and NbSPN14-2T7 (Table S1). A DsRNA-EGFP fragment was used as the mock group[42]. The amplified product was used as a template to synthesize dsRNA using a RiboMAX Large Scale System-T7 Kit (Promega, Madison, WI, USA). The dsRNA was then isolated and purified, stored at −80 °C. To evaluate the effect of RNAi, the ds-RNA and X-tremeGENE HP DNA Transfection Reagent were mixed at a ratio of 1:1 (m/v) and added dropwise to the BmE cells, which were then infected with *N. bombycis*. Host cell apoptosis was analyzed at 1dpi, 3dpi and 5dpi.

To block the expression of NbSPN14in silkworms, 10 μL (3 μg) dsRNA was injected into the silkworm hemocoel of the fifth instar; EGFP gene dsRNA was injected into the hemocoel of the control insects. These injections were performed immediately after the silkworms were orally inoculated with *N. bombycis* spores. A second injection of the same dose was administered after 24 hours. To confirm the interference efficiency, NbSPN14 transcript levels were determined by qRT-PCR.

### Yeast two-hybrid analysis

A yeast two-hybrid assay was used to investigate the interaction of NbSPN14 with BmICE. The detailed procedure was described previously (Bouzahzah et al., 2010). NbSPN14 was cloned into a pGADT7 plasmid, and PPAE was cloned into a pGBKT7 plasmid (Clontech, Takara Bio USA). The plasmids were used to transform competent yeast cells using YeastMaker Yeast Transformation System 2 (Takara, Mountain View, CA94043, USA), and the binding was validated in synthetic dropout-Leu-Trp-His-Ade medium supplemented with X-α-gal. The fusion strain of pGBKT7-53 with pGADT7-T was used as the positive control; the fusion strain of pGBKT7-lam with pGADT7-T was used as the negative control.

### Co-Immunoprecipitation (Co-IP)

Protein A + G agarose beads (Bio-Rad) were bound to V5 or HA antibody for 30 min, washed 3 times with 0.1% PBST, incubated with protein that had been extracted from NbSPN14-V5 and BmICE-HA recombinant protein co-expressed cells at 4 °C 4h, and then washed 3 times. The beads were then added (60 μL) to 1 × SDS PAGE loading buffer, the samples were boiled (10 min), and then used for western blotting.

### Construction BmICE knockout cell lines

sgRNA was designed based on the *BmICE* gene sequence and the CRISPRdirect online website (http://crispr.dbcls.jp/). Two sgRNA were designed in the fourth and fifth exon regions of the *BmICE* gene. Design primers based on the *BmICE* gene target sequence for expressing the sgRNA sequence fragment, then anneal the forward and reverse primers to form double-stranded DNA with sticky ends, which is connected to the sgRNA transcription vector pSL1180-U6-sgRNA containing the U6 promoter. 5’ end of the first gRNA target of the BmICE gene, and 3’ end downstream of the second gRNA target, 1000∼1500bp were selected respectively as the homologous arm sequence of the donor. The 5’ donor fragment, [oPIE1-*GFP::BleoR*-Sv40] fluorescence reporter resistance gene fragment, and the 3’ donor fragment were ligated into a pESI vector using a Hieff Clone® Plus Multi One Step Cloning Kit (Yisheng, 10912ES10, China). BmE cells were inoculated into a 6-well cell culture plate. Then, the pSL1180-U6-sgRNA vector and pESI donor vector were co-transfected into these BmE cells, and 72 hours after transfection, the screening medium containing 200 μg / mL Zeocin was replaced. The nutrient medium was replaced every 2 days until the green fluorescent cells ratio reached 98%.

### RNA extraction and qRT–PCR

The infected BmE cells and NbSPN14 transgenic cells’ total RNA was isolated using TRIzol regent (Invitrogen), and the mRNA in this preparation was then reverse transcribed into cDNA using the Hifair® AdvanceFast One-step RT-gDNA Digestion SuperMix for qPCR (Yeasen) according to the manufacturer’s instructions. qRT–PCR was then performed using the Hieff qPCR SYBR Green Master Mix (Yeasen), and *Nb-β tubulin* and *BmRPL3* were used as the internal control. Three biological replicates were performed for each experiment. All PCR primers used are listed in supplemental Table S1.

### Statistics

The difference between control and experimental assays was evaluated using a two-tailed Student’s T test employing GraphPad Prism 5 software. Statistical differences between two groups of p < 0.01 were selected to indicate highly significant differences, while p < 0.05 represented significant differences, and p ≥ 0.05 indicated the lack of significant difference.

## Acknowledgments

We are very grateful to Professor Louis M. Weiss from the Department of Pathology, Albert Einstein College of Medicine, for his guidance and assistance in writing the manuscript of the article.

---

**Fig. S1.**
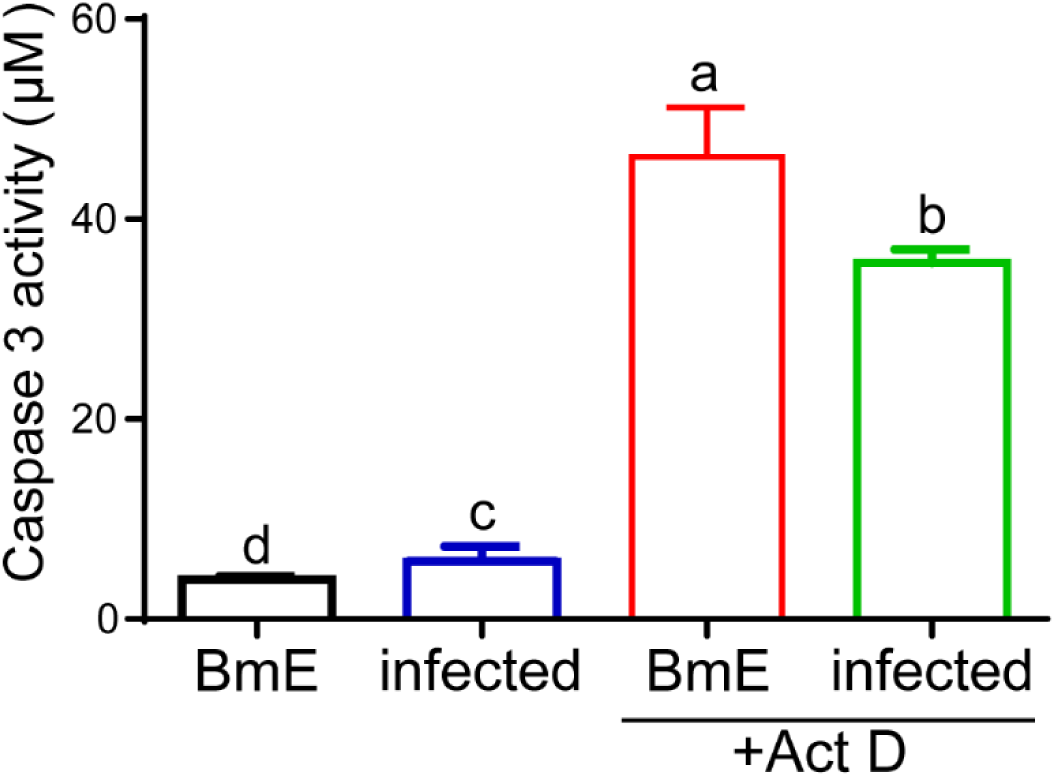
Analysis of Caspase 3 activity after *N. bombycis* infection and Act D induced apoptosis of BmE cells. The Caspase 3 activity of BmE cells infected with *N. bombycis* for 48 hours and treatment with Act D for 6 hours, Different and same letters indicate values with statistically significant (p < 0.05) and non-significant (p > 0.05) differences, respectively.

**Fig. S2.**
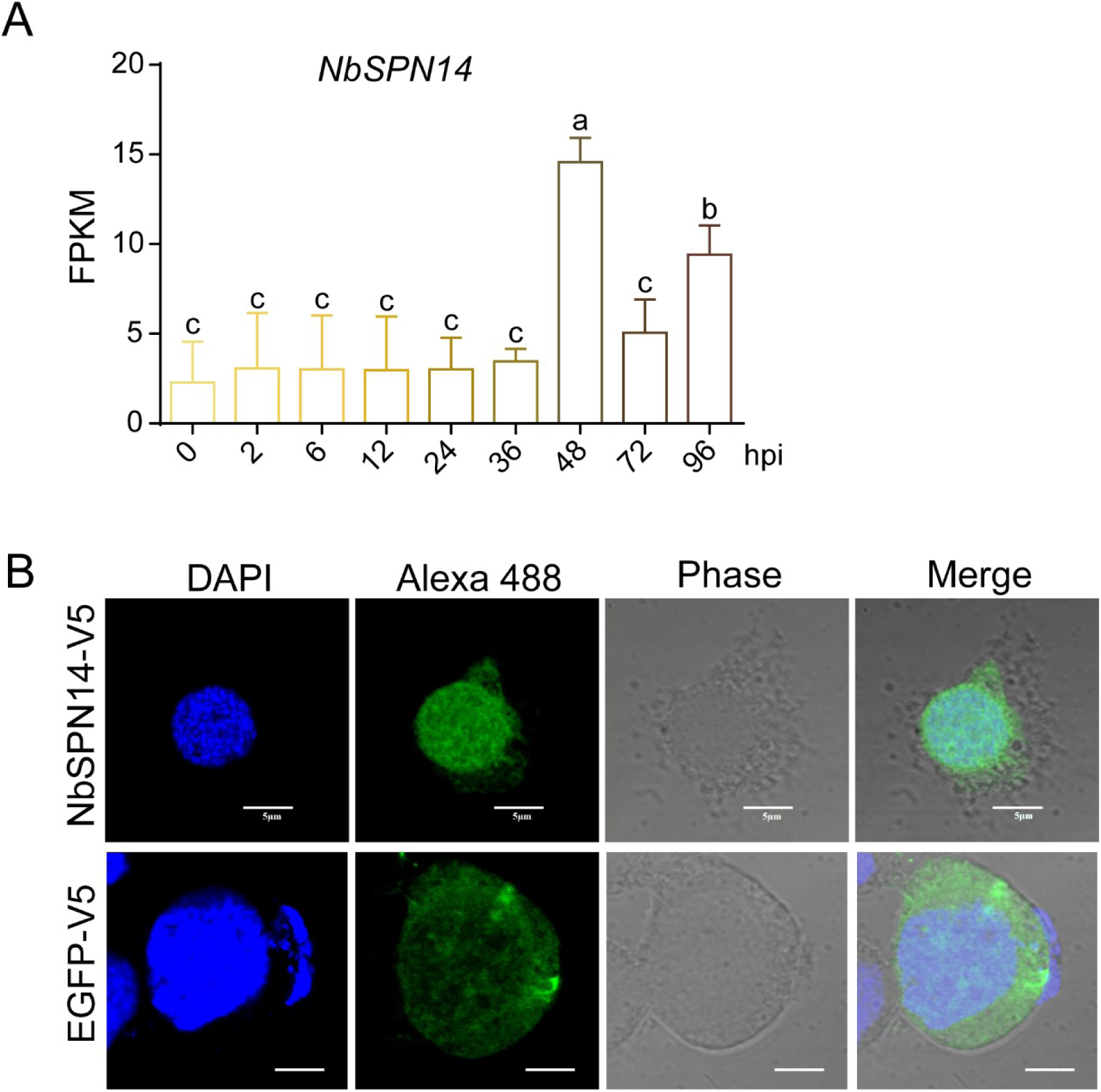
Transcriptional characterization of NbSPN14 in infected cell and expression localization of NbSPN14 in BmE cell. A. Analysis of NbSPN14 transcripts at 0, 2, 6, 12, 24, 36, 48, 72, and 92 h after infection with BmE cells showed that all transcripts were recorded, which was the highest at 48 h, followed by 96 h after infection. Different and same letters indicate values with statistically significant (p < 0.05) and non-significant (p > 0.05) differences, respectively. B. Forty-eight hours after transfection of the psl1180-IE2-NbSPN14-V5 expression vector into BmE cells, the localization of NbSPN14 fusion protein was analyzed by IFA. The primary antibody uses the V5-tagged mouse monoclonal antibody, the secondary antibody was the Goat anti-Mouse IgG with Alexa Fluor® 488.

**Fig. S3.**
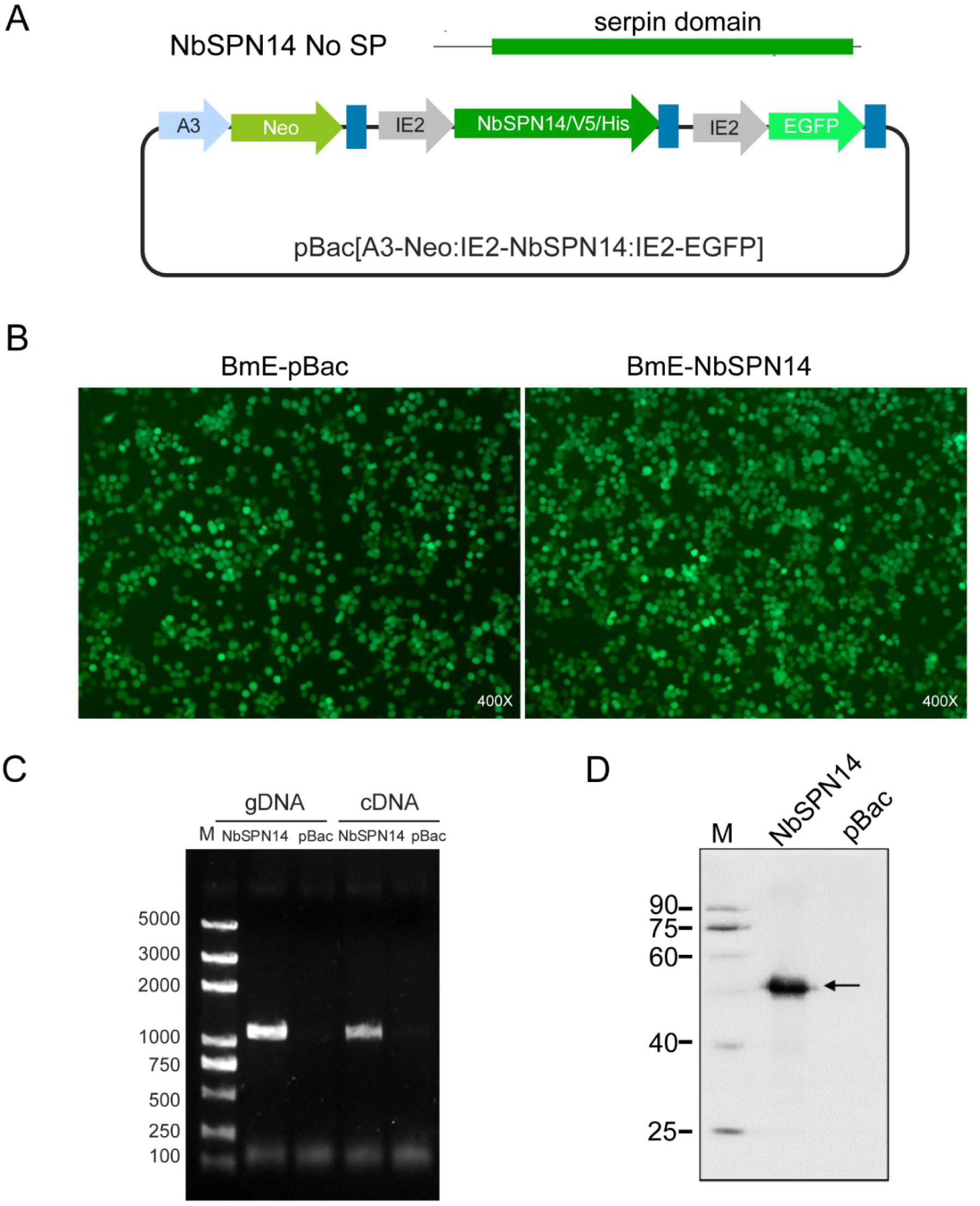
Construction and identification of NbSPN14 transgenic cells. A. The NbSPN14 signal peptide sequence was removed to mimic the secretion of NbSPN14 by *N. bombycis* into the host cell, and the schematic diagram of recombinant vector pBac [A3-Neo-SV40]-[IE2-NbSPN14-V5-SV40]-[IE2-EGFP-SV40], the vector pBac [A3-Neo-SV40] -[IE2-EGFP-SV40] as control; B. The pBac recombinant vector was transfected into BmE cells, and screened by Geneticin for 1 month. The percentage of cells with green fluorescence was more than 98%. C. The genomic DNA of transgenic cells was extracted, and the NbSPN14 gene fragment was successfully integrated into the host cell genome confirmed by PCR using a specific primer. The total RNA was extracted and reverse-transcribed into cDNA, and the successful transcription of NbSPN14 was verified. The pBac transgenic cells as a control. D. Total proteins from transgenic cells were extracted using RIAP lysate buffer. Western blotting was used to verify the expression of NbSPN14 in the transgenic cells;

**Fig.S4.**
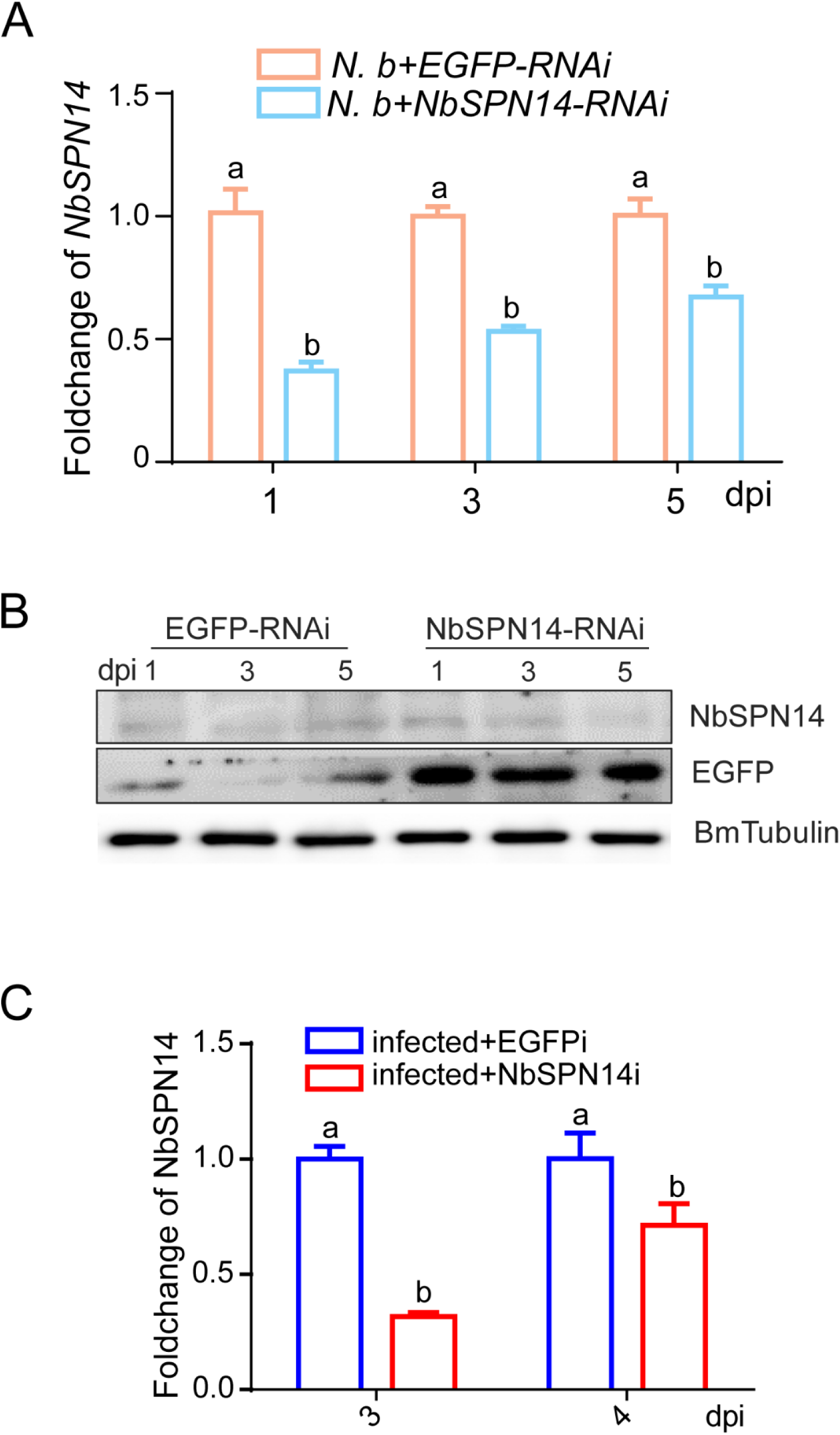
In *vitro* and *vivo* interference NbSPN14 expression verification. A. At the cellular level, NbSPN14 transcription analysis was performed after transfection of NbSPN14 interfered with double-stranded fragments, with transfected EGFP interference fragments as control. B. At the cellular level, Western blot was used to analyze the expression of NbSPN14 after transfection of interference fragments, and the transfected EGFP interference fragments were used as control. C. At the individual level, the transcription of NbSPN14 in the midgut was analyzed 3-4 days after injection of interference fragments into the stomata of silkworms infected with *N. bombycis*. Different and same letters indicate values with statistically significant (p < 0.05) and non-significant (p > 0.05) differences, respectively.

**Fig. S5.**
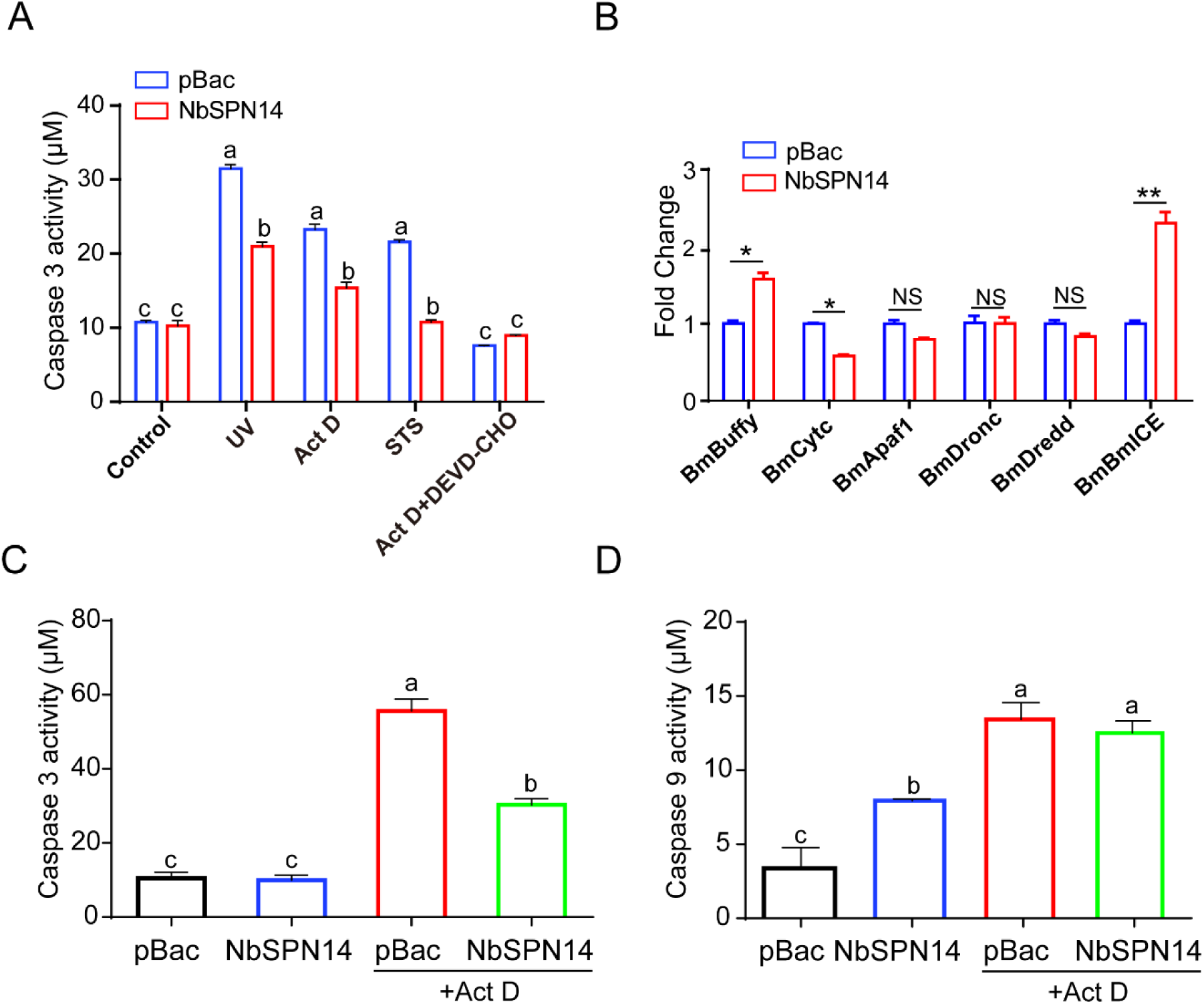
Analysis of NbSPN14 inhibition of apoptosis pathway in the host cell. A. NbSPN14 transgenic cells inhibit host cell Caspase 3 activity induced by Ultraviolet, Staurosporine (STS), and Actinomycin D (Act D) treatment. Ac-DEVD-CHO is a Caspase 3 specific inhibitor, as positive control. B. Quantitative PCR analysis was used to analyze the transcription of *BmBuffy*, *BmCytc*, *BmApaf1*, *BmDronc*, *BmDredd,* and *BmICE* genes related to the apoptosis pathway in NbSPN14 transgenic cells.; **indicating extremely significant difference (P < 0.01). C. Caspase 9 activity analysis of NbSPN14 transgenic cells with Act D treated 12h. D. Caspase 3 activity analysis of NbSPN14 transgenic cells treated with Act D for 12 h. Caspase 3 and Caspase 9 activity were assayed from the same sample, respectively. Different and same letters indicate values with statistically significant (p < 0.05) and non-significant (p > 0.05) differences, respectively.

**Fig. S6.**
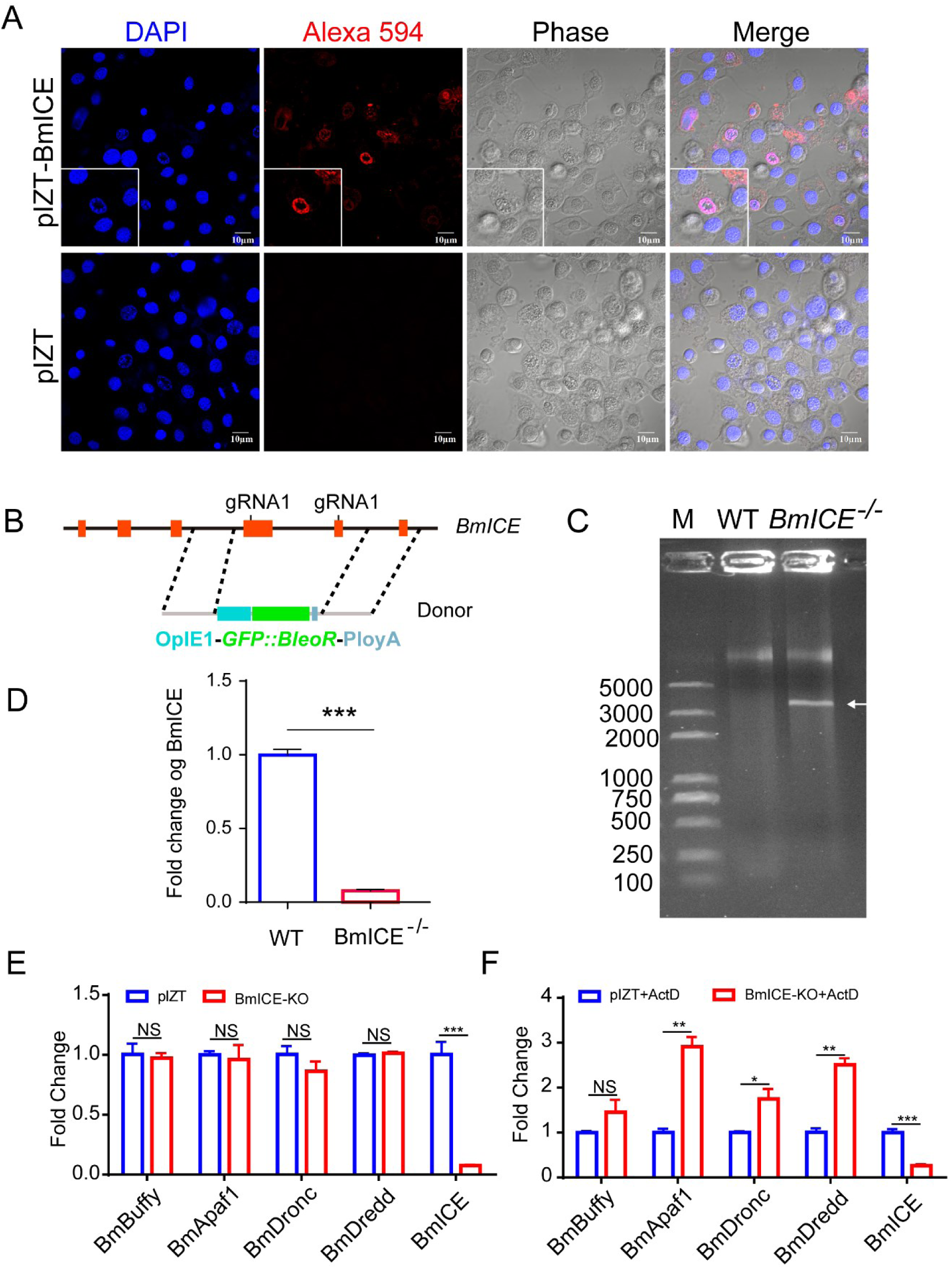
BmICE plays an important role in silkworm apoptosis. A. The PIZT-[IE1-BmICE-HA-sv40] expression vector was constructed and transfected into BmE cells. BmICE-HA is able to localize to the host cytoplasm and nucleus. Subcellular localization analysis of BmICE-HA recombinant proteins using HA-tagged rabbit antibody. The Alexa594-conjugated goat anti-rabbit antibody is the secondary antibody. B. Schematic diagram of the construction of BmICE knockout cells with two gRNA and homologous replacement fragment [OpIE1-*GFP::BleoR*-PloyA]; C. PCR verifies the integration of the gene fragment of the homologous donor [OpIE1-*GFP::BleoR*-PloyA] of the “GFP:: BleoR ” fragment into the genome. D. RT-qPCR analyzed the expression of *BmICE* in knockout cells. E. In *BmICE* knockout cells, RT-qPCR analysis of the *BmBuffy*/*BmApaf1*/*BmDronc*/*BmDredd*/*BmICE* gene expression in the apoptosis pathway showed that there was no significant difference between the experimental and the control group, only the expression of BmICE was significantly down-regulated. F. After the *BmICE* knockout cells were treated with Act D for 12 hours, the apoptosis-related genes *BmBuffy*/*BmApaf1*/*BmDronc*/*BmDredd*/*BmICE* genes expression was analyzed by quantitative PCR. The results showed that the apoptosis pathway was activated, but BmICE was still at a low transcriptional level.

